# Oct4-mediated inhibition of Lsd1 activity promotes the active and primed state of pluripotency enhancers

**DOI:** 10.1101/672451

**Authors:** Lama AlAbdi, Debapriya Saha, Ming He, Mohd Saleem Dar, Sagar M. Utturkar, Putu Ayu Sudyanti, Stephen McCune, Brice H. Spears, James A. Breedlove, Nadia A. Lanman, Humaira Gowher

## Abstract

An aberrant increase in pluripotency gene (PpG) expression due to enhancer reactivation could induce stemness and enhance tumorigenicity of cancer stem cells. Silencing of PpG enhancers (PpGe) during embryonic stem cell differentiation involves Lsd1–mediated H3K4me1 demethylation and DNA methylation. Here, we observed retention of H3K4me1 and DNA hypomethylation at PpGe associated with a partial repression of PpGs in F9 embryonal carcinoma cells (ECCs) post-differentiation. H3K4me1 demethylation in F9 ECCs could not be rescued by Lsd1 overexpression. Given our observation that H3K4me1 demethylation is accompanied by strong Oct4 repression in P19 ECCs, we tested if Oct4 interaction with Lsd1 affects its catalytic activity. Our data show a dose-dependent inhibition of Lsd1 activity by Oct4 and retention of H3K4me1 at PpGe in Oct4 overexpressing P19 ECCs. These data suggest that Lsd1-Oct4 interaction in cancer stem cells could establish a *primed* enhancer state that is susceptible to reactivation leading to aberrant PpG expression.

## Introduction

Cell type specific gene expression is regulated by chromatin conformation which facilitates the interaction of distally placed enhancer elements with the specific gene promoter (Banerji et al., 1981, Bulger and Groudine, 2011, Ong and Corces, 2011, Plank and Dean, 2014). Enhancers house the majority of transcription factor binding sites and amplify basal transcription, thus playing a critical role in signal dependent transcriptional responses (summarized in (Heinz et al., 2015)). Epigenome profiling combined with the transcriptional activity in various cell types led to identification of potential enhancers, which are annotated as silent, primed, or active based on their epigenetic features. These epigenetic features include histone modifications, and DNA methylation (Ernst and Kellis, 2010, Ernst et al., 2011, Calo and Wysocka, 2013). Whereas histone H3K4me1 (monomethylation) and H3K4me2 (dimethylation) is present at both active and primed enhancers, active enhancers invariantly are marked by histone H3K27Ac (acetylation) and/or transcribed to produced enhancer RNA (eRNA) (Heintzman et al., 2007, Heinz et al., 2010, Rada-Iglesias et al., 2011, Creyghton et al., 2010, Zentner et al., 2011, Zhu et al., 2013b).

During embryonic stem cell (ESC) differentiation, pluripotency gene (PpG) specific enhancers are silenced via changes in histone modifications and a gain of DNA methylation (Whyte et al., 2012, Mendenhall et al., 2013, Petell et al., 2016). In response to the differentiation signal, the coactivator complex (Oct4, Sox2, Nanog and Mediator complex) dissociates from the enhancer followed by the activation of pre-bound Lsd1-Mi2/NuRD enzymes. The histone demethylase Lsd1 demethylates H3K4me1, and the HDAC activity of the NuRD complex deacetylates H3K27Ac (Whyte et al., 2012). Our previous studies have shown that the histone demethylation event is critical for the activation of DNA methyltransferase Dnmt3a, which interacts with the demethylated histone H3 tails through its chromatin-interacting ADD (ATRX-Dnmt3a-Dnmt3L) domain, allowing site-specific methylation at pluripotency gene enhancers (PpGe) (Petell et al., 2016). These findings were further supported by biochemical studies showing that Dnmt3a-ADD domain interacts with histone H3 tail and this interaction is inhibited by H3K4 methylation (Guo et al., 2015, Li et al., 2011a, Ooi et al., 2007, Otani et al., 2009), together suggesting that aberrant inhibition of Lsd1 demethylase activity could cause a failure to gain DNA methylation, leading to incomplete repression of PpGs.

Several studies have reported on potential mechanisms that control site specific targeting and catalytic activity of Lsd1. Whereas Lsd1 interaction with CoREST activates the enzyme, BHC80 inhibits Lsd1 demethylation activity (Shi et al., 2005). The substrate specificity of Lsd1 is regulated by its interaction with androgen receptor and estrogen related receptor α, or by alternative splicing which adds four or eight amino acids to the Lsd1 enzyme (Carnesecchi et al., 2017, Metzger et al., 2005, Laurent et al., 2015, Zibetti et al., 2010, Wang et al., 2015a). Lsd1 is targeted to various genomic regions through its interaction with SNAG domain containing TFs, such as Snail and GFI1B (McClellan et al., 2019, Vinyard et al., 2019). The SNAG domain binds to the active site of Lsd1 by mimicking the histone H3 tail and could potentially inhibit its activity (Baron et al., 2011). Interaction of the p53 C’ terminal domain with the Lsd1 active site inhibits Lsd1 enzymatic activity (Speranzini et al., 2017). Lsd1 was also shown to be present in the Oct4 interaction network, and therefore could be targeted to Oct4-bound regulatory elements, which largely control pluripotency and stemness (van den Berg et al., 2010, Pardo et al., 2010, Ding et al., 2012).

Studies by the Cancer Genome Anatomy Project (CGAP) show that one out of three cancers express PpGs, suggesting their role in dysregulated proliferation during tumorigenesis (Zhang et al., 2013, Liu et al., 2013). Further, expression of PpGs, *Oct4*, *Sox2,* and *Nanog* potentiates self-renewal of putative cancer stem cells (CSCs) (Ben-Porath et al., 2008, Feske, 2007, Linn et al., 2010, Peng et al., 2010, Kumar et al., 2012, Wang et al., 2013, Mak et al., 2012, Wen et al., 2010, Jeter et al., 2011). CSCs proliferate as well as differentiate to give rise to cancer cells of various lineages (Iglesias et al., 2017). However, in order to retain the ability to proliferate, many cancer cells maintain expression of PpGs (Gwak et al., 2017, Yang et al., 2018). This has led to the development of terminal differentiation therapy, which aims to limit the proliferating cancer cell population (de The, 2018). Embryonal Carcinoma Cells (ECCs) have been used as a model cell line to study CSCs. ECCs were derived from developing mouse embryos at E6-7.5 and share regulatory characteristics with ESCs, including their ability to differentiate into various somatic lineages (Alonso et al., 1991, Han et al., 2017, Andrews et al., 2005, Zhu et al., 2013a). To understand the mechanism by which cancer cells retain PpG expression, we investigated the mechanism of enhancer-mediated regulation of PpG expression in ECCs. Our data showed that in differentiating F9 ECCs, the PpGs are only partially repressed. This was concomitant with H3K27 deacetylation, but with an absence of Lsd1-mediated H3K4me1 demethylation at PpGe.

The presence of H3K4me1 prevented Dnmt3a from methylating the DNA at these sites, potentially abrogating PpGe silencing. Drug-mediated inhibition as well as overexpression of Lsd1 had little or no effect on enhancer silencing and PpG repression, confirming an absence of Lsd1 dependence in differentiating ECCs. Given that Oct4 was expressed at substantial levels in F9 ECCs post-differentiation, we investigated the effect of Lsd1-Oct4 interaction on Lsd1 catalytic activity. Using *in vitro* histone demethylation assays, we discovered that Lsd1-Oct4 interaction inhibits Lsd1 activity, which could potentially result in the retention of H3K4me1 at PpGe in F9 ECCs. We tested this prediction in P19 ECCs in which Oct4 expression is strongly reduced post-differentiation and H3K4me1 is demethylated at PpGe. The observation that overexpression of Oct4 in differentiating P19 ECCs led to retention of H3K4me1 at PpGe, confirmed the role of Oct4-mediated Lsd1 inhibition at these sites. Taken together our data show that inhibition of Lsd1 activity and Dnmt3a leads to the establishment of a “primed” enhancer state, which is open for coactivator binding and prone to reactivation. We speculate that aberrant expression of Oct4 in CSCs facilitates the establishment of “primed” enhancers, reactivation of which support tumorigenicity.

## Results

### PpGs are partially repressed in differentiating F9 ECCs

ECCs share many characteristics with ESCs, including mechanisms governing regulation of gene expression and differentiation (Alonso et al., 1991). Based on the observation that aberrant PpG expression is commonly found in cancers (Zhang et al., 2013, Liu et al., 2013), we compared the magnitude of PpG repression in F9 ECCs with that in ESCs pre- and post-differentiation. ESCs and F9 ECCs were induced to differentiate with retinoic acid (RA) and expression of a subset of PpGs at 4 days (D4) post-induction was measured by RT-qPCR. In ESCs, the expression of most PpGs was reduced by more than 80% post-differentiation. The expression of *Sox2* and *Trim28* was maintained as an anticipated response to RA signaling guiding ESCs towards neural lineage (Figures 1A). However, in differentiating F9 ECCs, several PpGs were incompletely repressed, of which genes encoding the pioneer factors Oct4 showed 75% loss of expression whilst Nanog remain unchanged. A substantial increase in the expression of the genes *Lefty1* and *Lefty2* in F9 ECCs suggests potential transient activation of germ cell and testis developmental programs (Zhu et al., 2013a) (Figure 1B). To ensure that incomplete repression of PpGs was not the consequence of delayed response to RA signaling, we cultured cells for 8 days (D8). A variable decrease in PpGs was observed in the range of 25% - 75% with highest repression in *Lefty 2* (Figure 1B). Continued expression of PpGs in F9 ECCs was also evident by positive alkaline phosphatase staining and SSEA-1 immunofluorescence in differentiating F9 ECCs that is completely lost in ESCs post-differentiation (Figures 1C and 1D). We asked if failure to exit pluripotency in F9 ECCs was caused by an inability to activate lineage-specific genes. Our data showed a 5 to 60-fold increase in the expression of the lineage specific genes *Gata4, Foxa2, Olig2, Gata6, Cxcr4, and Fgf5* (Figure 1E), reflecting a standard response to differentiation signal. Given that previous studies in ESCs have established a critical role of enhancer silencing incomplete PpG repression, we next investigated if PpGe were fully decommissioned in F9 ECCs post-differentiation (Petell et al., 2016).

**Figure 1.**
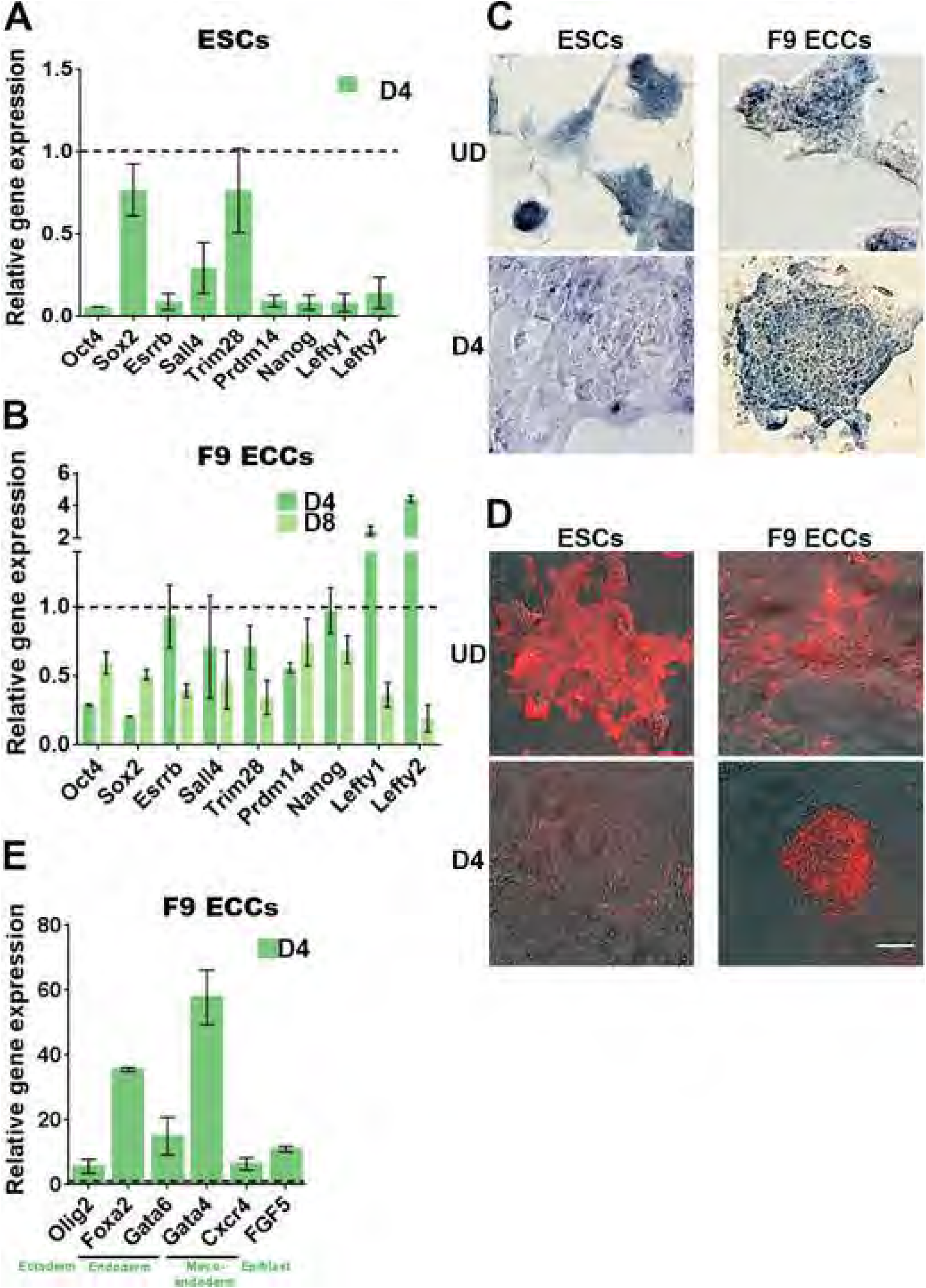
Pluripotency genes are partially repressed in embryonal carcinoma cells. UD: undifferentiated; D4, D8: Days post-induction of differentiation; ESCs: embryonic stem cells and F9 ECCs: F9 embryonal carcinoma cells, PpGs: pluripotency genes. (A, B, and E) Gene expression analysis by RT-qPCR of PpGs in (A) F9 ESCs, (B) ECCs (E) lineage specific genes in F9 ECCs. The C_t_ values for each gene were normalized to *Gapdh* and expression is shown relative to that in undifferentiated cells (dotted line). In F9 ECCs, the lineage specific genes show a 5 to 60-fold induction of gene expression (E) whereas the expression of PpGs is on average reduced to about 50% post-differentiation (B). Average and SEM of two biological replicates are shown for each gene. (C) Alkaline phosphatase staining and (D) SSEA-1 immunofluorescence of ESCs and F9 ECCs pre- and post-differentiation. Positive signal indicates pluripotency that is lost post-differentiation in ESCs. The scale bar is a 100 µm.

### DNA methylation is not established at PpGe during F9 ECC differentiation

In differentiating ESCs, PpGe silencing involves gain of DNA methylation, which is required for complete PpG repression (Petell et al., 2016). We used bisulfite sequencing (Bis-Seq) to compare DNA methylation changes at a subset of PpGe in F9 ECCs to that in ESCs post-differentiation. Whereas DNA methylation was significantly increased at most PpGe in ESCs 4 days post-differentiation, these sites remained hypomethylated (<10% methylation, (Tierling et al., 2018)) in F9 ECCs (Figures 2A and 2B). A similar hypomethylated state persisted at PpG promoters in F9 ECCs except the highly methylated *Lefty2* promoter, where DNA methylation was partially lost post-differentiation (Figure S1A). This result is consistent with the observed partial repression of most PpGs and an induction of *Lefty2* expression in these cells (Figure 1B). Furthermore, in ESCs, gain of DNA methylation at *Trim28* and *Sox2* enhancers suggests enhancer-switching, which involves a potential use of neural lineage specific enhancers to maintain a high expression of these genes post-differentiation (Figure 1A and 2A).

**Figure 2.**
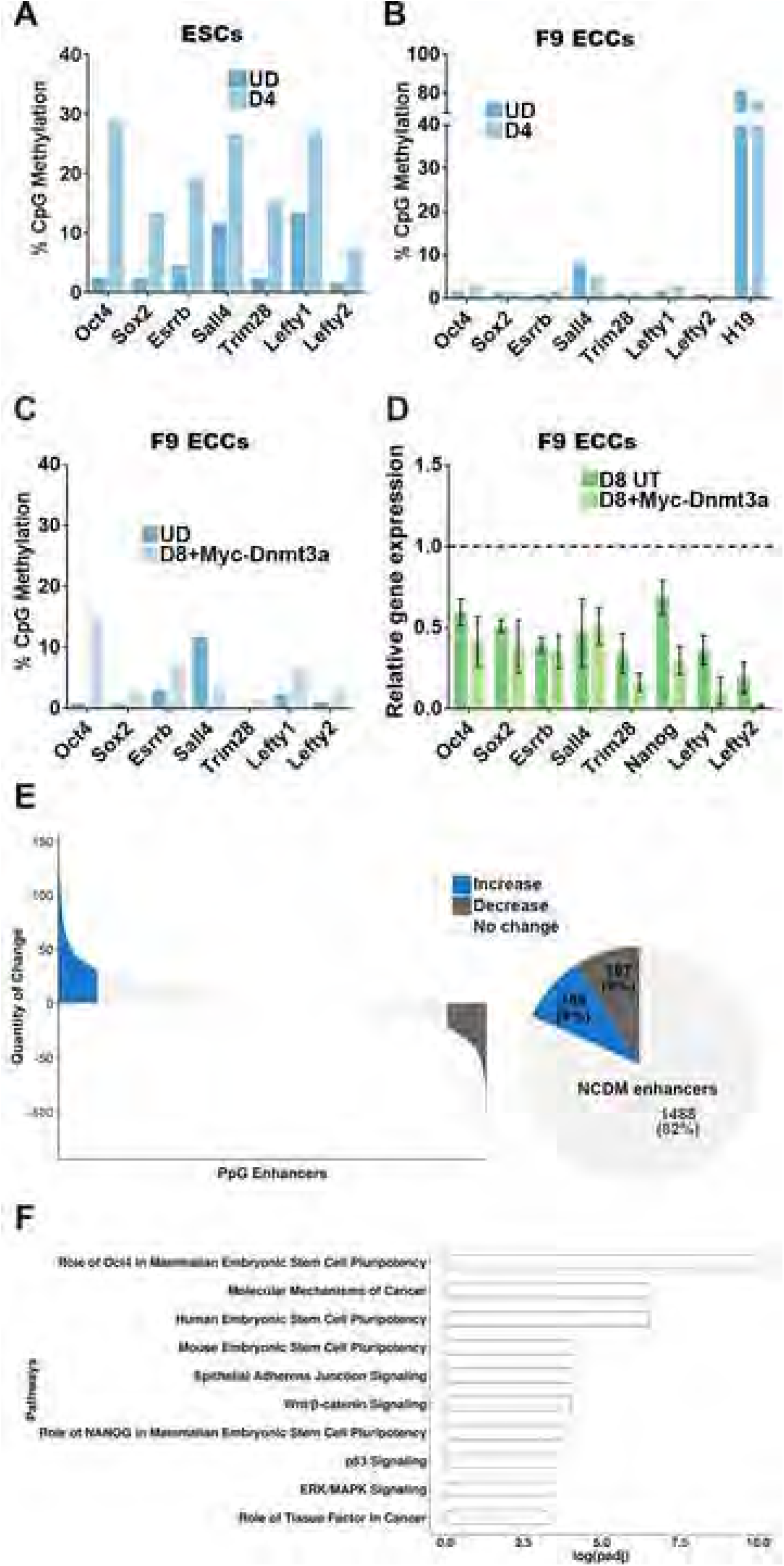
Pluripotency gene enhancers do not gain DNA methylation in embryonal carcinoma cells. UD: undifferentiated; D4, D8: Days post-induction of differentiation; D8 UT: untransfected F9 ECCs differentiated for 8 days; D8+Myc-Dnmt3a: F9 ECCs overexpressing Myc-Dnmt3a and differentiated for 8 days; ESCs: embryonic stem cells and F9 ECCs: F9 embryonal carcinoma cells; PpGe: pluripotency gene enhancers. (A, B, and C) DNA methylation analysis using Bis-Seq. Genomic DNA was treated with bisulfite and PpGe regions were amplified by PCR. The amplicons were sequenced on a high throughput sequencing platform (Wide-Seq) and the data were analyzed using Bismark software. DNA methylation of PpGe in (A) ESCs (B) F9 ECCs pre- and post-differentiation. Less than 10% DNA methylation was recorded in F9 ECCs whereas the H19 imprinted region, used as a control, showed DNA methylation at 80%. At the same regions, DNA methylation increased up to 30% in ESCs. See also Figure S1A. (C) DNA methylation of PpGe in F9 ECCs overexpressing Myc-Dnmt3a. (D) Gene expression analysis by RT-qPCR PpGs in F9 ECCs overexpressing Myc-Dnmt3a pre- and post-differentiation. (C) show low levels (less than 10%) in DNA methylation at most PpGe (D) no significant decrease in expression in 5 out of 8 tested PpGs (*P*-value >0.1). The C_t_ values for each gene were normalized to *Gapdh* and expression is shown relative to that in undifferentiated cells (dotted line). Data for A, B, C and D are the average and SEM of two biological replicates. (E) Genome-wide DNA methylation analysis by MethylRAD sequencing. Genomic DNA was digested with the restriction enzyme *Fsp*EI that cuts methylated DNA into 31-32 bp fragments. The fragments were sequenced and mapped on mm10 mouse genome. The number of reads per region were used as a measure for extent of DNA methylation and compared between undifferentiated and D4 differentiated F9 ECCs. The waterfall plot shows DNA methylation changes at PpGe computed by subtracting normalized counts in D4 samples from normalized counts in undifferentiated samples. Upper and lower quartiles were used in thresholding regions as gaining or losing methylation. The pie chart shows fractions of PpGe with increase, decrease or no change in DNA methylation (NCDM). See also Figure S2B. (F) Top ten statistically significant enriched canonical pathways amongst the genes associated with the NCDM enhancers, which showed no change. The x-axis shows the log_10_ (adjusted p-value), with the p-value adjusted for multiple testing using the Benjamini-Hochberg method.

We confirmed that absence of DNA methylation at PpGe was not due to low expression of Dnmt3a in F9 ECCs post-differentiation (Figures S1B and S1C). Based on previous observations in cancers that overexpression of DNA methyltransferases leads to DNA hypermethylation (Yu et al., 2015b, Gao et al., 2015, Ma et al., 2018, Jones et al., 2016, Schübeler, 2015), we tested if overexpression of Dnmt3a could rescue DNA methylation at PpGe. F9 ECCs were transfected with Myc-Dnmt3a and differentiated at 24 hr post-transfection to ensure expression of recombinant Dnmt3a during early differentiation (Figure S1D). We anticipate the DNA methylation established by transiently over expressing Dnmt3a will be maintained by Dnmt1 during multiple cell divisions (Lyko, 2018). To ensure the detection of methylation established by recombinant Dnmt3a, we differentiated the cells for 8 days. However, DNA methylation levels at PpGe except Oct4 (15%) were well below <10% which is within the range of detection error by this method (Tierling et al., 2018). A small gain within this range was observed suggesting a spurious low methylation during cell divisions (Figure 2C). 5 out of 8 PpGs showed no additional decrease in expression (*p* value >0.1) when compared to untransfected differentiated cells (Figure 2D) indicating that overexpression of Dnmt3a is unable to rescue the differentiation defects observed in F9 ECCs. The absence of complete PpG repression concomitant with little or no significant gain in DNA methylation at PpGe predicts a potential disruption in the mechanism that mediates PpGe decommissioning.

To determine the extent of DNA methylation defect, we perfromed MethylRAD sequencing to analyze changes in DNA methylation at all PpGe in F9 ECCs pre- and post-differentiation (Wang et al., 2015b). This method uses *Fsp*EI restriction enzyme, which cuts DNA bidirectionally from mC to create 31-32 bp fragments (Cohen-Karni et al., 2011, Zheng et al., 2010). The restriction fragments were isolated for library preparation and high throughput sequencing. Using this method, we captured DNA methylation at 1,370,254 cytosines genome-wide. The reads were distributed among all chromosomes representing all annotated genomic elements (Figure S2A). DNA methylation levels at enhancers (low-intermediate-high) were calculated based on highest (75th percentile) and lowest (25th percentile) number of reads at all annotated enhancers in the genome, which were obtained from the EnhancerAtlas 2.0. Previous studies reported that PpGe are bound by Lsd1 in ESCs and post-differentiation, the PpGe decommissioning requires H3K4me1 demethylation by Lsd1 (Whyte et al., 2012). In succession, our studies showed that H3K4me1 demethylation was critical for Dnmt3a catalyzed DNA methylation at these sites (Petell et al., 2016). Therefore, we filtered the data to focus our analysis on DNA methylation changes at 3840 PpGe previously annotated in ESCs as Lsd1 bound regions (Whyte et al., 2012). Our method identified 1,865 PpGe in F9 ECCs. Compared to methylation levels at all other known enhancers, the PpGe cluster into the low/intermediate methylation group (Figure S2B). The difference in methylation for each PpGe region was computed by subtracting the DNA methylation level in D4 differentiated from that in undifferentiated F9 ECCs. The data show 1488 (82%) regions fail to gain DNA methylation post-differentiation of F9 ECCs (No Change in DNA Methylation; NCDM) (Figure 2E). To determine the function of genes associated with NCDM PpGe, we performed *Ingenuity* pathway analysis (IPA) (www.qiagen.com/ingenuity), which showed a significant enrichment of Oct4-regulated mammalian embryonic stem cell and molecular mechanisms of cancer pathways (Figure 2F).

Given that DNA methylation by Dnmt3a at PpGe requires H3K27 deacetylation and H3K4 demethylation by Lsd1/Mi2/NurD complex (Whyte et al., 2012), we anticipate a potential impediment in this process causing a widespread failure to acquire DNA methylation at PpGe.

### A “primed” PpGe state is established during F9 ECC differentiation

We asked if the chromatin state at PpGe in F9 ECCs is refractory to DNA methylation. To examine histone H3K27 deacetylation, chromatin immunoprecipitation followed by qPCR (ChIP-qPCR) was performed that showed a decrease in H3K27Ac at PpGe in F9 ECCs post-differentiation (Figure 3A). This result suggests that, similar to our observations in ESCs (Figure S3A), PpGe are active in undifferentiated F9 ECCs and initiate the decommissioning process post-differentiation. Furthermore, deacetylation at *Lefty1* and *Lefty2* enhancers suggests enhancer-switching involving the potential use of germline-specific enhancers post-differentiation, leads to an observed increase in *Lefty1* and *Lefty2* expression (Figure 1A). Numerous studies have proposed that deacetylation of H3K27Ac followed by H3K27 methylation by the PRC2 enzyme complex establishes a silenced state (Lindroth et al., 2008, Barski et al., 2007, Wang et al., 2008). Our data showed no increase of H3K27me3 at the PpGe in both ESCs as well as F9 ECCs post-differentiation, suggesting that PRC2 activity is nonessential for PpGe silencing (Figures S3B and S3C). H3 occupancy at the PpGe in F9 ECCs between pre- and post-differentiation was similar, supporting these conclusions (Figure S3D).

**Figure 3.**
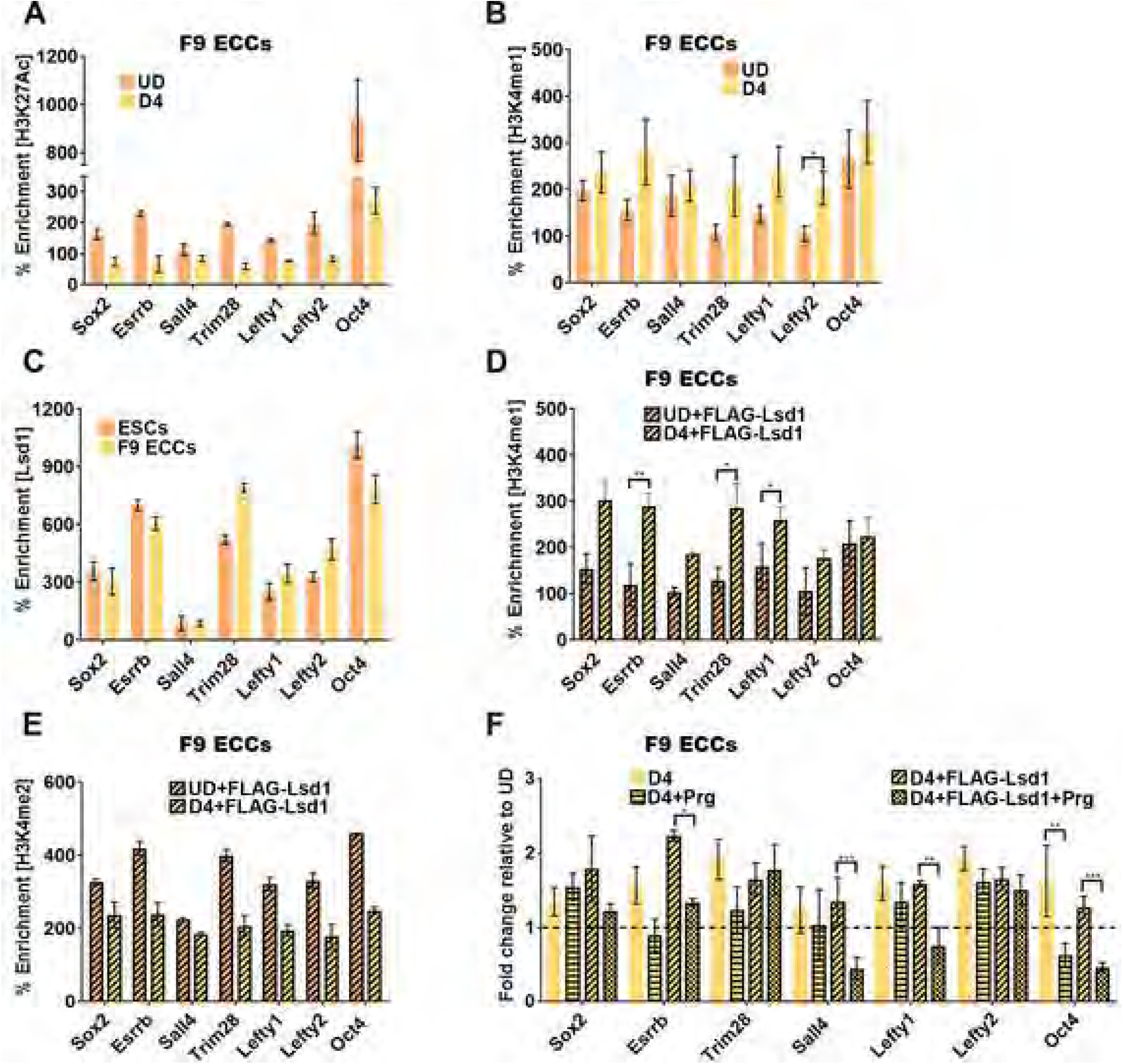
A “primed” chromatin state is established at pluripotency gene enhancers in embryonal carcinoma cells. UD: undifferentiated; D4: Days post-induction of differentiation; D4+FLAG-Lsd1: F9 ECCs overexpressing FLAG-Lsd1 and differentiated for 4 days; Prg: Pargyline; ESCs: embryonic stem cells and F9 ECCs: F9 embryonal carcinoma cells; PpGe: pluripotency gene enhancers. (A-F) Chromatin immunoprecipitation (ChIP)-qPCR assays. Histone modifications at PpGe (A) H3K27Ac and (B) H3K4me1 in F9 ECCs pre- and D4 post-differentiation. Whereas deacetylation of PpGe is observed as a decrease in the H3K27Ac signal, histone H3K4me1 is retained post-differentiation. (C) Lsd1 occupancy in undifferentiated ESCs and F9 ECCs. (D, E) Enrichment of (D) H3K4me1 and (E) H3K4me2 in F9 ECCs expressing recombinant FLAG-Lsd1 compared to undifferentiated transfected cells. The ECCs were differentiated 24hrs post transfection with Lsd1 expressing plasmid. Whereas there is an increase in H3K4me1 at some PpGe (D), there is a concomitant decrease in H3K4me2 enrichment (E). (F) Fold change in enrichment of H3K4me1 at PpGe in pargyline treated and untreated, WT, and FLAG-Lsd1 overexpressing F9 ECCs, at D4 post-differentiation. Fold change is represented as relative to enrichment in the undifferentiated state (dotted line). % Enrichment = Fold enrichment over input X100.*P*-values derived from Student’s *t*-test where *, **, and *** denote (*P*-value <0.05), (*P*-value <0.01), and (*P*-value <0.005), respectively.

Using ChIP-qPCR, we next monitored H3K4me1 demethylation at PpGe during F9 ECC differentiation. Surprisingly, we observed a retention of H3K4me1 modification at most enhancers and a significant increase at 1 out of 7 tested PpGe post-differentiation (Figure 3B), suggesting a potential disruption of Lsd1 activity. We verified similar expression levels of Lsd1 in F9 ECCs compared to ESCs (Figures S1B and S1C). Previous studies have shown Lsd1 to interact with the components of NuRD complex in ESCs and cancer cells to facilitate deacetylation of H3K27 followed by demethylation of H3K4me1 (Whyte et al., 2012, Petell et al., 2016, Li et al., 2011b, Patel et al., 2018, Wang et al., 2009b). To examine if Lsd1 interacts with the Mi2/NuRD complex in F9 ECCs, we performed co-immunoprecipitation (Co-IP) experiments using whole cell extracts. We used the whole cell extract from ESCs as a positive control. Antibodies against Lsd1 and HDAC1 were used for reciprocal Co-IP. A strong signal for HDAC1 and Lsd1 and a weak signal for CHD4 in both F9 ECCs and ESCs was observed (Figure S3E). An absence of signal for the acetyltransferase HBO1 in Lsd1 Co-IP and presence of CHD4 supports the specificity of the interaction between Lsd1 and NuRD complex in F9 ECCs (Figure S3F). To test whether Lsd1 is recruited to PpGe, ChIP-qPCR showed similar enrichment of Lsd1 in F9 ECCs and ESCs at PpGe (Figure 3C). These data suggest that retention of H3K4me1 at PpGe is not caused by the absence of Lsd1 but rather due to lack of its activity at these sites in F9 ECCs post-differentiation.

To confirm the above conclusion, we next tested the effect of overexpression or inhibition of Lsd1 on PpGe silencing and PpG repression in differentiating F9 ECCs. F9 ECCs were transfected with a FLAG-Lsd1 overexpressing plasmid and differentiated at 24 hr post-transfection (Figure S4A). The recombinant Lsd1 was not able to rescue H3K4me1 demethylation at PpGe shown by no decrease in H3K4me1 at most PpGe (Figure 3D). However, a significant increase was observed at 3 out of 7 PpGe post-differentiation. This could result from incomplete demethylation of H3K4me2 to H3K4me1 at these sites by recombinant Lsd1, confirmed by a decrease in H3K4me2 signal at most of the PpGe in Lsd1 overexpressing F9 ECCs post-differentiation (Figure 3E). These data suggest that an inhibitory mechanism affects Lsd1 demethylation activity irrespective of its origin of expression. Additionally, Lsd1 overexpression had no significant effect on PpG repression or DNA methylation at PpGe post-differentiation (Figures S4B and S4C).

Previous studies in ESCs have shown that treatment with the Lsd1 inhibitor pargyline at the onset of differentiation results in H3K4me1 retention at PpGe and incomplete repression of PpGs (Whyte et al., 2012). F9 ECCs (WT and Lsd1 overexpressing) were treated with pargyline 6 hr prior to induction of differentiation. In contrast to 70-80% cell death caused by the Lsd1 inhibitor in ESCs, WT F9 ECCs remained largely viable upon treatment (Figure S4D) and a slight relief of repression was observed for some PpGs in these cells (Figure S4E). Whereas no decrease in H3K4me1 was observed at most PpGe in WT F9 ECCs, pargyline treatment affected the H3K4me1 enrichment at 4 out of 7 PpGe in Lsd1 overexpressing cells post-differentiation. This confirms the activity of Lsd1 at these sites by which it converts H3K4me2 to H3K4me1 (Figure 3F). To test if Lsd1 activity contributed to the generation of H3K4me1 at PpGe in the undifferentiated state, we transiently overexpressed Lsd1 in F9 ECCs and allowed cells to grow for 72 hr. ChIP-qPCR analysis showed no increase in H3K4me1 levels in Lsd1 overexpressing cells (Figure S4F). This result suggests that, similar to what has been previously reported in ESCs, the deposition of H3K4me1 in the undifferentiated F9 ECCs is largely accomplished by MLL3/4 histone methyltransferases, and Lsd1 activity at these sites is initiated only in response to a differentiation signal (Wang et al., 2017, Wang et al., 2016, Cao et al., 2018, Whyte et al., 2012).

Taken together, these observations suggest that the restricted activity of Lsd1 at PpGe leads to retention of H3K4me1 post-differentiation. Moreover, following the deacetylation of H3K27, the absence of DNA methylation and the presence of H3K4me1 switches these enhancers to a “primed” state, prone to reactivation.

### High-throughput analysis of changes in H3K4me1 at PpGe

To identify all PpGe with aberrant retention of H3K4me1 post-differentiation, we performed ChIP-Seq analysis of H3K4me1 genome-wide in undifferentiated and D4 differentiated F9 ECCs. Peak calling was performed using Epic2 for each input-ChIP pair and the list was filtered to calculate the number of peaks based on defined cut offs (FDR<=0.05 and Log2FC>=2) (Figure S5A). We analyzed the distribution of H3K4me1 peaks at the regulatory elements across the genome (Figure S5B). For further analysis, we identified 1425 H3K4me1 peaks found within 1 kbp of PpGe previously annotated in ESCs (Whyte et al., 2012). The difference in peak enrichment (i.e. log2FoldChange) between differentiated and undifferentiated was calculated to score for changes in H3K4me1 post-differentiation. The data showed no change, increase, or decrease in 733, 510, and 182 PpGe, respectively. Therefore 87% of PpGe showed no significant decrease in H3K4me1 (no decrease in histone methylation; NDHM PpGe) (Figure 4A). We next computed the correlation between the three PpGe sub-groups and the 1792 PpGe that undergo H3K4me1 demethylation in differentiating ESCs (Whyte et al., 2012). Among the NDHM PpGe, 74%PpGe with increased H3K4me1 and 69% PpGe with unchanged H3K4me1 in F9 ECCs, overlap with PpGe that are H3K4me1 demethylated in ESCs (Figures 4B, 4C, and S5C). These observations strongly support our previous conclusion that in F9 ECCs, Lsd1 activity is inhibited leading to retention of H3K4me1 at PpGe. IPA of NDHM PpGe–associated genes showed highest enrichment for Oct4-regulated and stem cell pathways (Figure 4D). Comparatively, genes associated with PpGe that undergo H3K4me1 demethylation, show enrichment for signaling pathways (Figure S5D). A correlation between NCDM and NDHM PpGe showed that 65% of the NDHM PpGe fail to acquire DNA methylation, underpinning the role of histone demethylation in the regulation of DNA methylation at PpGe (Figure 4E) (Petell et al., 2016).

**Figure 4.**
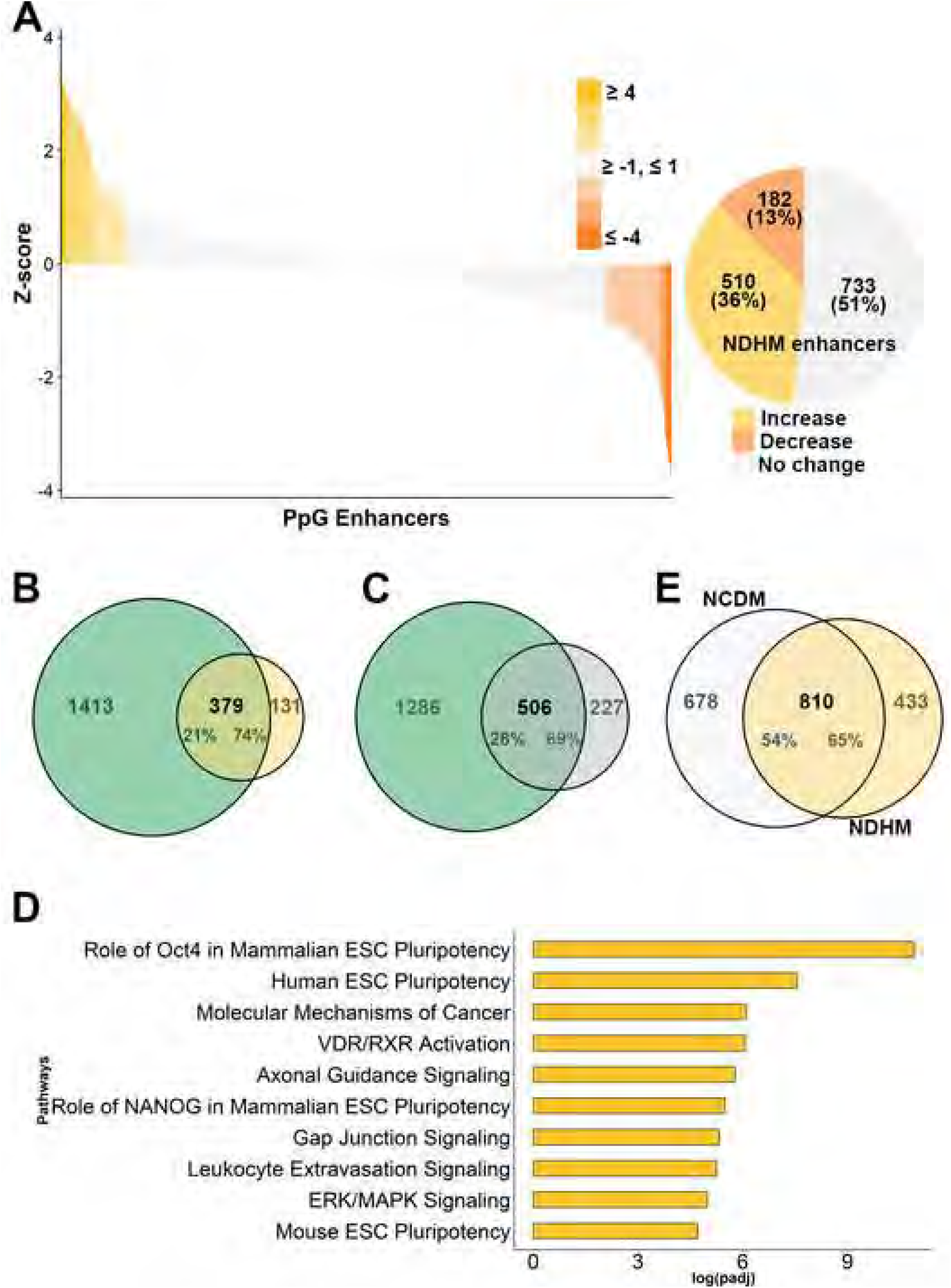
Global retention of H3K4me1 at pluripotency gene enhancers in embryonal carcinoma cells. ESCs: embryonic stem cells and F9 ECCs: F9 embryonal carcinoma cells; PpGe: pluripotency gene enhancers; D4: Days post-induction of differentiation Genome-wide H3K4me1 levels in F9 ECCs pre- and post-differentiation were measured by ChIP-Seq. Peak calling was performed using Epic2 for each input-ChIP pair. A total of 1425 H3K4me1 peaks were identified in F9 ECCs within 1 kbp of previously annotated PpGe in ESCs (Whyte et al., 2012). See also Figure S5A. Histone demethylation activity of Lsd1 was surmised by calculating the change in H3K4me1 peak enrichment at PpGe between D4 differentiated and undifferentiated samples. (A) Waterfall plot represents changes in H3K4me1, which were calculated as the difference between log2FC of D4 and undifferentiated samples, and transformed to Z-score. Z-score thresholds of +1 and −1 were used to define the fractions showing increase, no change, or decrease in H3K4me1 shown in the pie chart. Taken together, 87% PpGe show an increase or no change (NDCM) in H3K4me1 enrichment. (B, C) Venn diagram showing an overlap between PpGe that show (B) an increase, or (C) no change in peak enrichment in F9 ECCs but undergo histone H3K4me1 demethylation in ESCs post-differentiation (Whyte et al., 2012). (D) Top ten statistically significant enriched canonical pathways amongst the genes associated with increase and no change in F9 ECCs. The x-axis shows the log_10_ (adjusted p-value), with the p-value adjusted for multiple testing using the Benjamini-Hochberg method. (E) Overlap between the PpGe showing no change in DNA methylation (NDHM) and PpGe that show no decrease in H3K4me1 (NCDM).

### Lsd1 activity at PpGe is inhibited by its interaction with Oct4

Next, we sought to determine the mechanism that inhibits Lsd1 activity. Due to its continued expression in F9 ECCs post-differentiation (Figure S6A), we assumed that Oct4 remains associated with PpGe and prevents demethylation of H3K4me1 at these sites. We tested this hypothesis in P19 ECCs in which Oct4 expression was reported to be strongly repressed post-differentiation (Li et al., 2007, Wei et al., 2007, Li et al., 2013, Palmieri et al., 1994, Fuhrmann et al., 2001, Liu et al., 2011, Marikawa et al., 2011), an observation distinct from what we report in F9 ECCs. After confirming a 90% reduction in Oct4 expression (Figure 5A), we probed H3K4me1 demethylation at PpGe during P19 ECC differentiation. Indeed, our data show decreased enrichment of H3K4me1 at the 5 tested PpGe, 4 days post-differentiation (Figure 5B). The change in chromatin state included up to 40% gain of DNA methylation at 3 out of 5 enhancers (Figure 5C). Similar to ESCs, we also observed a massive cell death when P19 ECCs were exposed to the Lsd1 inhibitor during differentiation (Figure S6B).

**Figure 5.**
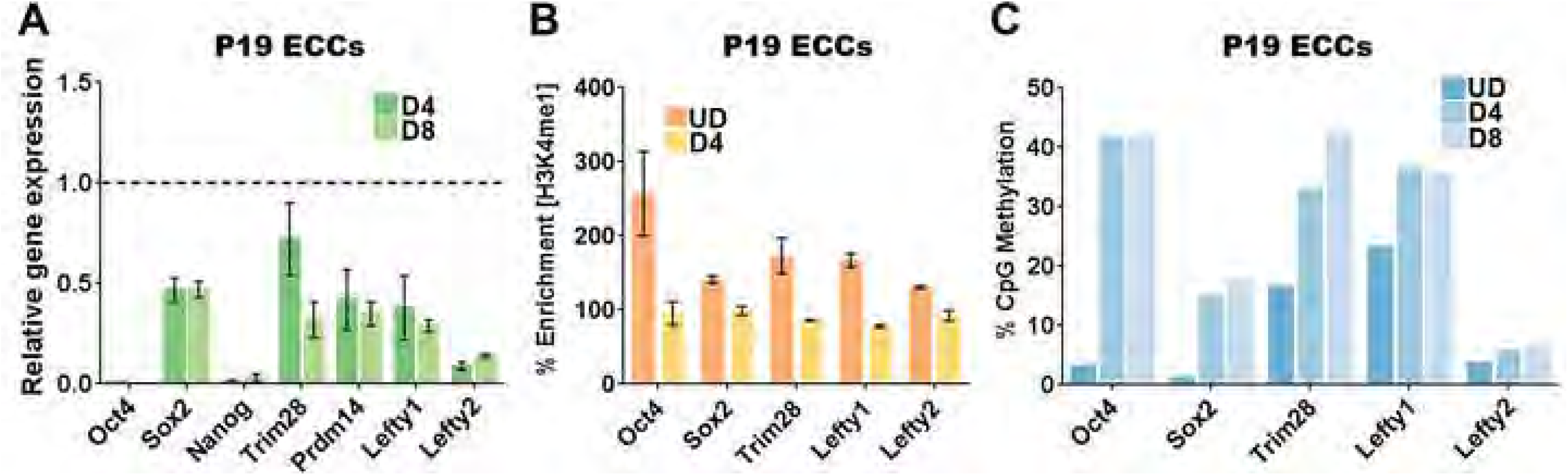
Pluripotency gene enhancers are decommissioned in P19 embryonal carcinoma cells. UD: undifferentiated; D4, D8: Days post-induction of differentiation; P19 ECCs: P19 embryonal carcinoma cells, PpGs: pluripotency genes; PpGe: Pluripotency gene enhancers. (A) Gene expression analysis by RT-qPCR of PpGs in P19 ECCs. The C_t_ values for each gene were normalized to *Gapdh* and expression is shown relative to that in undifferentiated cells (dotted line). Similar to the repression of PpGs in ESCs (Figure 1A), PpGs, especially Oct4 and Nanog, show more than 90% reduction in expression. (B) ChIP-qPCR showing H3K4me1 enrichment in UD and D4 differentiated P19 ECCs. A decrease in H3K4me1 was observed at all PpGe post-differentiation demonstrating histone demethylation activity. % Enrichment = Fold enrichment over input X100. (C) DNA methylation analysis of PpGe using Bis-Seq in UD, D4 and D8 differentiated P19 ECCs. Up to 40% increase in DNA methylation level was observed at 3 out of 5 PpGe post-differentiation. Average and SEM of two biological replicates are shown for each gene.

Together with the pathway analysis showing enrichment of Oct4-regulated genes in NDHM and NCDM enhancers, the above data suggested that Oct4 might regulate the demethylase activity of Lsd1 at PpGe. Previous observations showing an interaction of Oct4 with Lsd1 and the Mi2/NuRD complex support this assumption (van den Berg et al., 2010, Pardo et al., 2010, Ding et al., 2012). Interaction of Lsd1 and Oct4 was confirmed by co-immunoprecipitation of Lsd1 with anti-Oct4 antibody using nuclear extract from both undifferentiated and D4 differentiated F9 ECCs (Figure 6A). Co-precipitation experiments using recombinant proteins, GST-Lsd1 and Oct4, reveal a direct interaction between the two proteins (Figure 6B). To test the effect of Oct4 interaction on Lsd1 catalytic activity, we performed *in vitro* Lsd1 demethylation assays using H3K4me2 peptide as a substrate. In the presence of Oct4, Lsd1 activity was reduced by 60-70% in a dose dependent manner. The specificity of Oct4-mediated inhibition was demonstrated by no effect of recombinant Dnmt3a protein on Lsd1 activity. Complete loss of Lsd1 activity in the presence of its inhibitor TCP (tranylcypromine) confirmed the specificity of the signal measured in this assay (Figures 6C and 6D).

**Figure 6.**
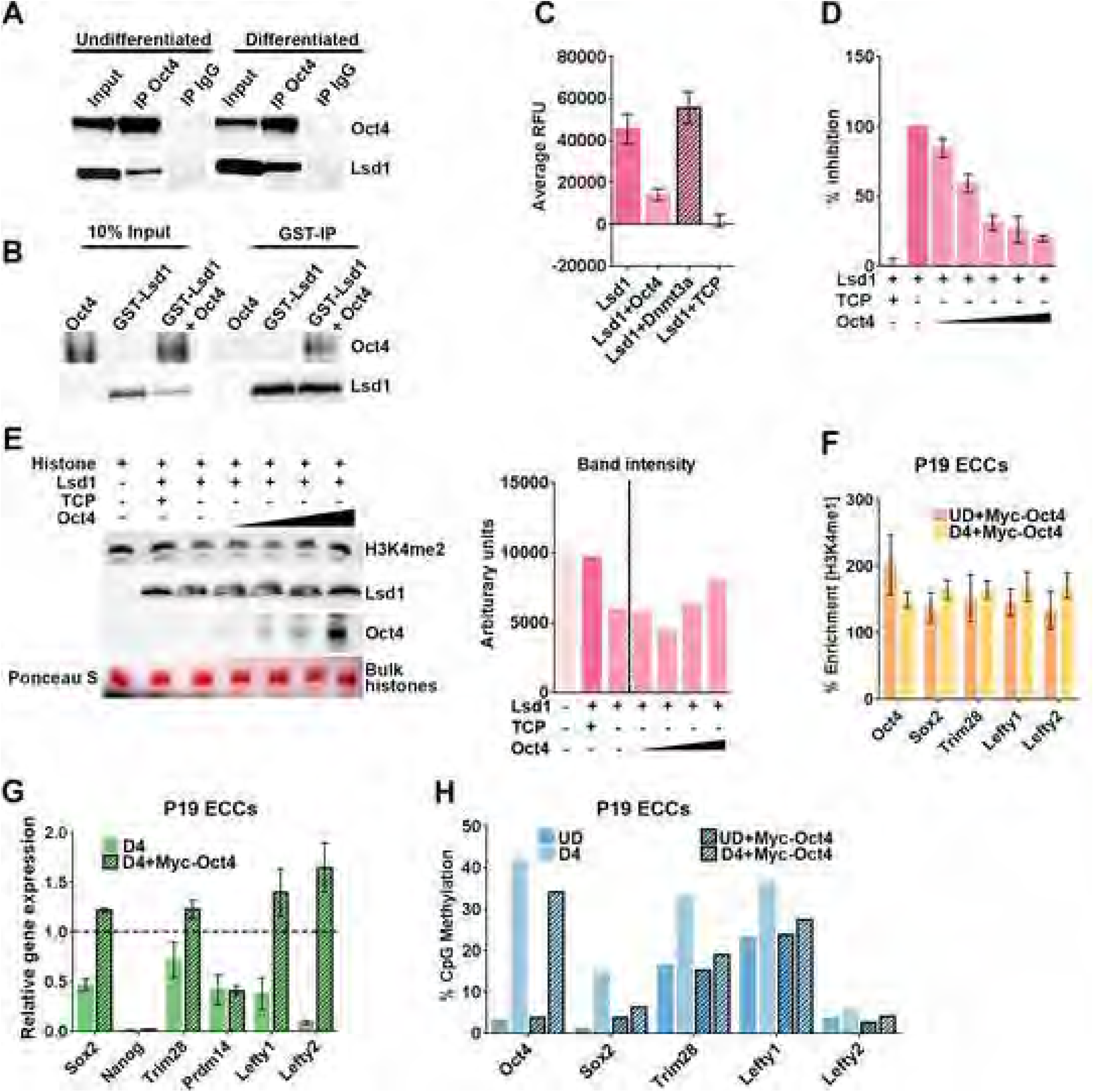
Oct4 interacts with Lsd1 and inhibits its catalytic activity. TCP: tranylcypromine; ECCs: embryonal carcinoma cells; PpGe: pluripotency gene enhancers D4: Days post-induction of differentiation. (A) Nuclear extract from undifferentiated and D4 differentiated F9 ECCs was used to perform Co-IP with anti-Oct4 antibody and control IgG. 10% of the input and eluate from Co-IP were probed for Oct4 and Lsd1 on Western blot. The vertical space denotes the extra lane in the gel that was digitally removed. (B) GST-pull down experiment showing direct interaction between Lsd1 and Oct4. Recombinant GST-Lsd1 was incubated with Oct4 at about 1:2 molar ratio and precipitated using GST-Sepharose. The co-precipitated Oct4 is detected using Anti-Oct4 antibody. The vertical space denotes the extra lane in the gel that was digitally removed (C) Lsd1 demethylase assay was performed using 0.25 µM of Lsd1 and H3K4me2 peptide as substrate. Lsd1 demethylation activity was completely inhibited by 0.1 mM TCP (tranylcypromine) in the reaction. To test the effect of Oct4 on Lsd1 activity, demethylation assays were performed in the presence of 0.5 µM Oct4 at 1:2 (Lsd1:Oct4) molar ratio. The catalytic domain of Dnmt3a at the same molar ratio was used as a control. (D) Dose dependent inhibition assays were performed using increasing concentrations of Oct4 in the following molar ratios of Lsd1:Oct4, (1:0.5), (1:1), (1:2), (1:3), (1:4). Data are an average and SD of at least 5 experimental replicates. (E) Lsd1 demethylation assays were performed using 0.25 µM of Lsd1 and 30 µg bulk histones as substrate with increasing concentrations of Oct4 in the reaction. Upper Panel: Histone demethylation was detected by using anti H3K4me2 on Western blot showing a retention of signal in presence of increasing concentration of Oct4. Lower panels: Amount of Lsd1 enzyme and increasing amounts of Oct4 in the histone demethylation reaction. Ponceau S stain of bulk histones shows equal loading on the gel. The bar graph on the right shows quantification of H3K4me2 signal using imageJ software. (F) ChIP-qPCR showing percent enrichment of H3K4me1 at PpGe in P19 ECCs stably expressing recombinant Myc-Oct4 pre- and post-differentiation. The data show retention of H3K4me1 post-differentiation. % Enrichment = Fold enrichment over input X100 (G) Gene expression analysis by RT-qPCR of PpGs in P19 ECCs expressing recombinant Myc-Oct4. The C_t_ values were normalized to *Gapdh* and expression is shown relative to that in undifferentiated cells (dotted line). (H) DNA methylation analysis of PpGe using Bis-Seq in UD and D4 differentiated P19 ECCs WT and expressing recombinant Myc-Oct4. Oct4 expressing cells show failure to gain DNA methylation at PpGe post-differentiation compared to untransfected WT. Average and SEM of two biological replicates are shown.

We also performed Lsd1 demethylation assays using purified histones as a substrate and detected H3K4me2 demethylation on a Western blot. Activity of Lsd1 was assessed by reduced H3K4me2 signal, which was rescued in the presence of 0.1 mM TCP. An accumulation of H3K4me2 signal with an increase in Oct4 concentration in the reaction mix clearly showed increased inhibition of Lsd1 activity by Oct4 (Figure 6E). These data suggest that Oct4 could inhibit Lsd1 activity at PpGe in ECCs post-differentiation. We tested this by stably expressing recombinant Oct4 in P19 ECCs (Figure S6C). Upon differentiation, the retention of H3K4me1 at PpGe indicates the inhibition of Lsd1 by the recombinant Oct4 (Figure 6F). Moreover, several PpGs were incompletely repressed with reduced gain in DNA methylation at their respective enhancers (Figures 6G and 6H). These data suggest that in F9 ECCs, due to its continued expression post-differentiation, Oct4 remains bound at PpGe and inhibits Lsd1 activity.

Taken together, we propose the following model to explain the regulation of Lsd1 activity at PpGe. At the active PpGe, Lsd1 is inhibited by its interaction with bound Oct4. Post-differentiation, this inhibition is relieved by dissociation of Oct4 from its binding sites. However, in cancer cells where Oct4 expression is maintained, Lsd1 is held in its inhibited state, leading to incomplete histone demethylation. Retention of H3K4me1 in turn blocks the activation of Dnmt3a from its autoinhibited state, resulting in an absence of DNA methylation at PpGe (Petell et al., 2016, Guo et al., 2015, Li et al., 2011a, Ooi et al., 2007, Otani et al., 2009). The absence of DNA methylation and the presence of H3K4me1 switches these enhancers to a “primed” state, prone to activation in the presence of the coactivator (Figure 7). Based on previous reports that Oct4 as well as Lsd1 are aberrantly expressed in several cancers (Kim et al., 2015, Wang et al., 2010, Schoenhals et al., 2009, Wang et al., 2009b, Kashyap et al., 2013, Lv et al., 2012, Hosseini and Minucci, 2017), we suspect that Oct4-Lsd1 interaction at Oct4-bound regions will disrupt Lsd1 activity leading to aberrant gene expression.

**Figure 7.**
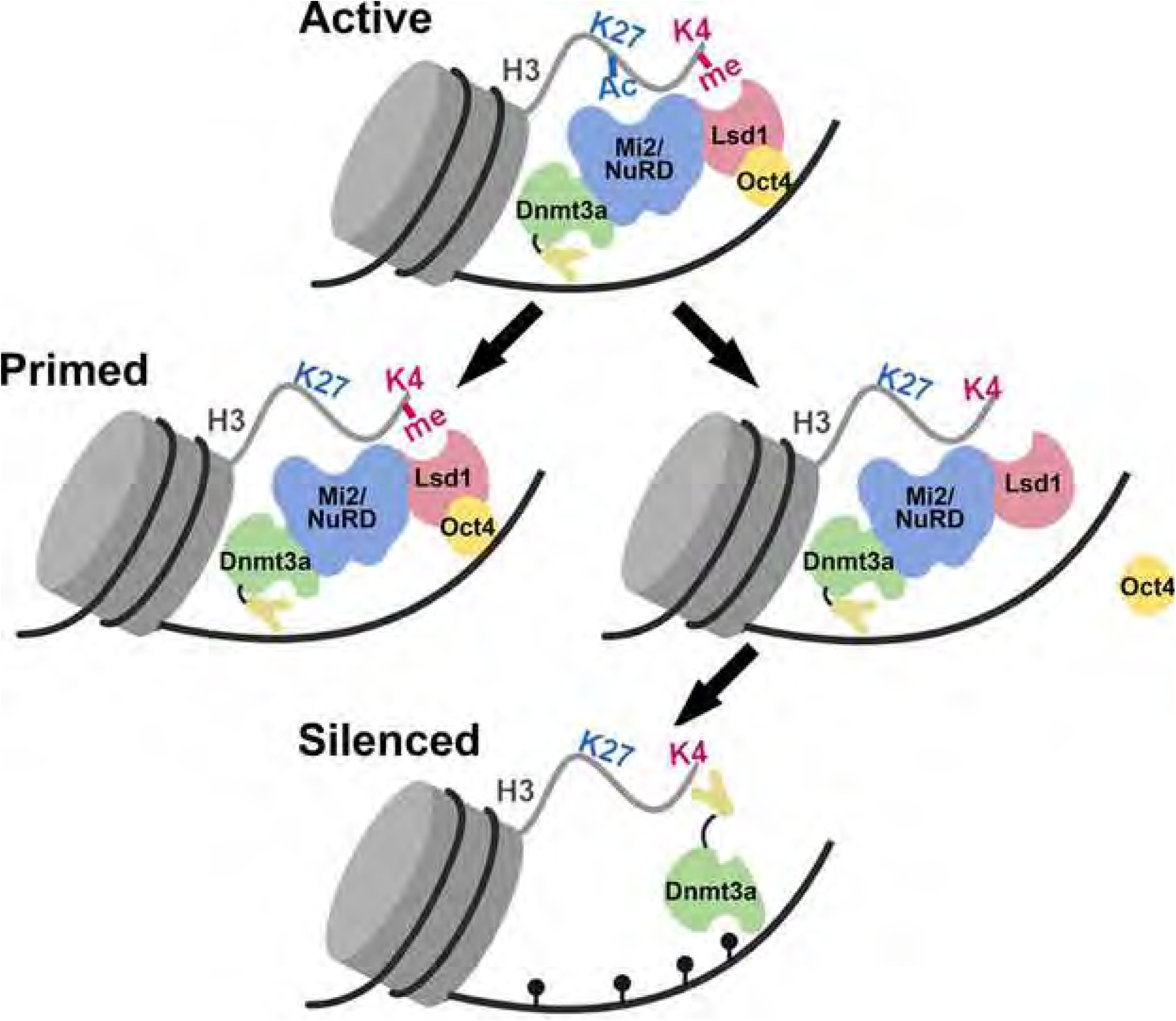
Model of epigenetic changes at pluripotency gene enhancers during stem cell differentiation. In an undifferentiated state, the pluripotency gene enhancers (PpGe) are active, bound by the coactivator complex and containing chromatin modifications including H3K4m2/1 and H3K27Ac. In response to signal of differentiation, the dissociation of the coactivator complex including Oct4 is followed by the activity of the Lsd1-Mi2/NuRD complex, which facilitates enhancer silencing. The histone deacetylase (HDAC) removes H3K27Ac at PpGe, and Lsd1 demethylates H3K4me1, followed by DNA methylation by Dnmt3a. However, in F9 ECCs, Lsd1 activity is inhibited in the presence of Oct4, causing retention of H3K4me1. The ADD domain of Dnmt3a cannot interact with H3K4 methylated histone tail and will potentially remain in the autoinhibited state, thus preventing DNA methylation at these sites. Consequently, PpGe instead of being silenced acquire a “primed” state. Black pins represent methylated CpGs.

## Discussion

The epigenetic state of enhancers is preserved in various cell types suggesting that aberrant changes could promote tumorigenesis. This phenomenon is supported by recent studies revealing a crucial role for enhancer-mediated activation of oncogenes (Hnisz et al., 2015, Mansour et al., 2014, Chapuy et al., 2013, Groschel et al., 2014, Loven et al., 2013). Changes in H3K4me1 levels and DNA accessibility at various enhancers have been reported in many cancers (Akhtar-Zaidi et al., 2012). Some studies also propose that changes in enhancer states play a role in the development of therapy resistant cancer cells. These studies showed loss or gain of H3K4me1/2 at the enhancers in resistant breast cancer cells and loss of H3K27 acetylation at enhancers in T cell acute lymphoblastic leukemia (T-ALL) (Magnani et al., 2013, Knoechel et al., 2014). In addition, DNA hypermethylation, concomitant with overexpression of DNA methyltransferases, is a hallmark of many cancers (Jones et al., 2016, Schübeler, 2015). Changes in DNA methylation at enhancers occur in breast, lung, prostate, and cervical cancers (Taberlay et al., 2014, Aran and Hellman, 2013, Aran et al., 2013, Yegnasubramanian et al., 2011). These studies suggest that the chromatin state of tissue-specific enhancers can be used as a diagnostic to predict aberrant expression of their respective genes in cancer. This prediction is supported by our data here and previous studies showing that PpGe gain DNA methylation and lose H3K4me1 in differentiating ESCs, whereas in ECCs, absence of DNA methylation is accompanied by retention of H3K4me1 at these sites. The tissue specificity of the enhancer state is further highlighted by our data showing that during ESC differentiation, all tested PpGe undergo repressive chromatin changes irrespective of the transcriptional status of the associated gene. This observation is exemplified by H3K27Ac deacetylation and gain of DNA methylation at *Sox2* and *Trim28* enhancers despite the maintained expression of these genes.

Histone demethylase, Lsd1, catalyzes demethylation of dimethyl (me2) and monomethyl (me1) at K4 of histone H3 (Shi et al., 2004). The catalytic activity of Lsd1 was shown to be regulated by post-translational modification, alternative splicing and interaction with several factors (Carnesecchi et al., 2017, Laurent et al., 2015, Metzger et al., 2005, Shi et al., 2005, Speranzini et al., 2017, Wang et al., 2015a, Zibetti et al., 2010, McClellan et al., 2019, Vinyard et al., 2019). Genetic knockout of the histone demethylase, Lsd1, was also shown to cause genome-wide loss of DNA methylation in late passage ESCs due to degradation of DNMT1 enzyme (Wang et al., 2009a). Our previous studies demonstrated the critical role of H3K4me1 demethylation by Lsd1 in guiding DNMT3A-mediated DNA methylation at PpGe causing enhancer silencing in ESCs (Petell et al., 2016). In contrary to ESCs, our data in differentiating F9 ECCs shows that PpGs are partially repressed and their respective enhancers retain H3K4me1 and DNA hypomethylation. The transgenic overexpression of Dnmt3A or Lsd1 in F9 ECCs was unable to rescue H3K4me1 demethylation and gain of DNA methylation at these PpGe. A small gain of DNA methylation (15%) was observed at the Oct4 enhancers in Dnmt3A –overexpressing cells, however this was insufficient to further downregulate the expression of Oct4. This could be explained by retention of histone H3K4me1 at the Oct4 enhancer that keeps the enhancer in a partially active state. Moreover we show that when Oct4 gene is repressed during P19 ECCs differentiation, Oct4 enhancer accumulates upto 40% DNA methylation accompanied by H3K4me1 is also demethylated (Figure 5). This suggest that higher DNA methylation levels are required for complete silencing of Oct4 enhancer. The data above is in agreement with previous studies showing a similar accumulation DNA methylation at the Oct4 enhancer (40%-50%) when Oct4 gene is repressed during ESCs post differentiation (Athanasiadou et al., 2010, Petell et al., 2016). Taken together our data that compromised activity of Lsd1 results in the retention of H3K4me1 and the absence of DNA methylation at the PpGe, leading to a primed state of enhancers in cancer cells. Thus we reveal a mechanism by which developmental enhancers could acquire aberrant histone modification and DNA methylation states that affect gene expression. Unlike the silenced state, the “primed” enhancer state grants accessibility for coactivator binding which renders cells highly vulnerable to a small increase in the expression of oncogenic coactivators or master transcription factors (Zaret and Carroll, 2011, Calo and Wysocka, 2013).

We also discovered that Lsd1 activity is inhibited by its interaction with the pioneer transcription factor Oct4, which is expressed at a substantial level in F9 ECCs post-differentiation. Recently, a similar observation was reported from flow cytometric analysis showing that compared to ESCs, a significantly higher number of F9 ECCs have persistent *Oct4* expression post-differentiation (Gordeeva and Khaydukov, 2017). Aberrant expression of *Oct4*, *Sox2*, and *Nanog* is associated with tumor transformation, metastasis, and drug resistance (Sampieri and Fodde, 2012, Ben-Porath et al., 2008). We speculate that during differentiation of cancer stem cells, inhibition of Lsd1 by Oct4 leads to PpGe priming/reactivation leading to enhanced PpG expression. Our studies further highlight the versatile regulation of Lsd1 binding and activity, which can be fine-tuned by its interaction with numerous factors, allowing the enzyme to function in various cellular processes including differentiation and disease.

## Acknowledgements

We are thankful to Gowher lab members for discussions. This work was supported by NIHR01GM118654-01 and graduate fellowship for LA from King Saud University. The authors gratefully acknowledge the Walter Cancer Foundation, DNA Sequencing Facility, and support from the Purdue University Center for Cancer Research, P30CA023168. We thank Dr. Phillip SanMiguel from Genomic Core, Purdue for analysis of Bis-Seq data and Dr. Taiping Chen for providing FLAG-Lsd1 and Myc-Dnmt3a2 expression plasmids.

## Authors Contributions

L.A., S.M., B.H., J.B., M.H., D.S., and M.S.D performed the experiments. N.A., S.U., and P.A.S. analyzed the genome-wide data. L.A. and H.G. wrote the manuscript

## STAR Methods

### LEAD CONTACT AND MATERIALS AVAILABILITY

Further information and requests for resources and reagents should be directed to and will be fulfilled by the Lead Contact, Humaira Gowher

All unique/stable reagents generated in this study will be made available on request but we may require a payment for processing and shipping and/or a completed Materials Transfer Agreement if there is potential for commercial application.

### EXPERIMENTAL MODEL AND SUBJECT DETAILS

F9 embryonal carcinoma cells (F9 ECCs), P19 embryonal carcinoma cells (P19 ECCs), and E14Tg2A Embryonic stem cells (ESCs) were cultured and maintained in gelatin-coated tissue culture plates. All three above cell lines are male. Differentiation of ECCs was induced by plating 20X10^6^ cells in low attachment 15 cm petri dishes and the addition of 1µM Retinoic acid (RA). ESCs were differentiated by the same method with a concurrent withdrawal of LIF. The medium was replenished every two days and samples were collected on Days 4 and 8 post-differentiation.

Plasmids expressing Myc-Dnmt3a2 and FLAG-Lsd1 WT were transfected into F9 ECCs using Lipofectamine 2000. One day post-transfection, a UD sample was collected (D0), and transfected cells were induced to differentiate on gelatinized plates by the addition of RA. The next day, differentiated cells were trypsinized and plated on low adherence petri dishes. Samples were collected on Days 4 and 8 post-differentiation. Lsd1 inhibitor treatment was performed as described 6 hr prior to induction of differentiation (Petell et al., 2016).

P19 ECCs were transfected with pCAG-Myc-Oct4 (Addgene 13460) using Lipofectamine 2000 per the manufacturer’s instructions. Transfected cells were clonally propagated, and Myc-Oct4 expression was determined by Western blots with anti-cMyc antibody (Millipore, MABE282).

### METHOD DETAILS

#### DNA methylation analysis

Bisulfite sequencing: Bisulfite conversion was performed using an EpiTect Fast Bisulfite Conversion Kit (Qiagen, 59802) according to the manufacturer’s protocol. PCR conditions for outer and inner amplifications were performed (Petell et al., 2016). The pooled samples were sequenced using NGS on a Wide-Seq platform. The reads were assembled and analyzed by Bismark and Bowtie2. Methylated and unmethylated CpGs for each target were quantified, averaged, and presented as percent CpG methylation. Number of CpGs for regions tested are listed in Table S1. Total number of reads used to calculate percent CpG methylation are listed in Table S2, S3. Primer sequences can be found in Table S4.

MethylRAD sequencing: Genomic DNA was isolated using a standard phenol:chloroform extraction, followed by ethanol precipitation. DNA from various samples was digested with *Fsp*EI for 4 hr at 37°C and subjected to electrophoresis through a 2% agarose gel. 30 base pair fragments were cut out, purified, and adaptors were ligated at 4°C overnight (Wang et al., 2015b). The ligated DNA was PCR amplified with index primers and sequenced using a Novaseq 6000. The primers used for PCR amplification is in Table S4. The details of the bioinformatics analysis of data are listed in Supplementary Methods.

#### Chromatin Immunoprecipitation and ChIP-Seq

ChIP was performed as described (Petell et al., 2016). Chromatin was sheared by sonication using a Covaris E210 device, according to the manufacturer’s protocol. A total of 8µg of sheared crosslinked chromatin was incubated with 8µg of antibody pre-loaded on a 1:1 ratio of protein A and protein G magnetic beads (Life Technologies, 10002D and 10004D, respectively). After washing the beads, the samples were eluted in 1% SDS, 10 mM EDTA, 50 mM Tris-HCl, pH 8.0. Crosslinking was reversed by incubation at 65°C for 30 min with shaking. Samples were treated with RNase (Roche, 11119915001) for 2 h at 37°C, and subsequently treated with Proteinase K (Worthington, LS004222) for 2 h at 55°C. DNA was purified by phenol:chloroform extraction followed by ethanol precipitation and quantified using PicoGreen (Life Technologies, P11495) and NanoDrop 3300 fluorospectrometer. qPCR was then performed using equal amounts of IN and IP samples. Fold enrichment was first calculated as: 2^(C_t_(IN)-C_t_(IP)). Percent enrichment = Fold enrichment X 100. Significance of change was determined via *p*-value, which was calculated by GraphPad Prism using Student’s *t*-test.

Table S4 lists sequences of primers used.

#### Gene expression analysis

RNA was isolated using the TRIzol reagent (Invitrogen, 15596026) according to the manufacturer’s protocol. Samples were treated with DNAse (Roche, 04716728001) at 37°C, and then purified using a Quick-RNA^TM^ MiniPrep Plus Kit (ZymoReseach, R1057). Reverse-transcription quantitative PCR was performed by using Verso One-Step RT-qPCR kits (Thermo Scientific, AB-4104A) with 1µg of purified RNA. Gene expression was calculated as ΔC_t_ which is C_t_(Gene)-C_t_(*Gapdh*). Change in gene expression is reported as fold change relative to that in undifferentiated cells, which was set to 1. See Table S4 for primers used.

#### Microscopy

Bright field images of embryoid bodies (EBs) were obtained with Zeiss microscope using a 10X objective. Alkaline phosphatase staining was performed using solutions supplied by an alkaline phosphatase staining kit (Sigma, AB0300). Cells were cross-linked with 1% formaldehyde for 5 min, followed by quenching with a final concentration of 150 mM glycine. Cells were washed twice with 1xPBS, then twice with combined staining solution (BCIP and NBT). The stain was developed in the dark for 5 min, then washed three times with 1XPBS. SSEA-1 immunofluorescence was performed using the following antibodies: anti-SSEA-1 (Millipore, MAB430) and AlexaFluor 555 nm (Life Technologies, A21422). SSEA-1 and Alkaline phosphatase staining were imaged using 20X objectives under Nikon Ts and Zeiss microscopes, respectively.

#### Co-precipitation assays

For co-immunoprecipitation (Co-IP), the nuclear extract was prepared according to manufacturer’s protocol (Active Motif, 40010) except that DNase was added for the release of chromatin-associated proteins. The Co-IP was performed using 1:1 mix of Dynabeads™ Protein A (Life Technologies, 1002D) and Dynabeads™ Protein G (Life Technologies, 1004D), and conjugated with 5 μg antibody and with 50 μg of nuclear extract according to the manufacturer’s protocol. Antibodies used include: anti-Oct4 (Abcam, ab181557), anti-CHD4 (Abcam, ab72418), anti-HDAC1 (Abcam, ab7028), anti-Lsd1 (Abcam, ab17721) and anti-HBO1 (Abcam, ab124993).

Pull down assays were performed using 1 µg of GST-Lsd1 (Sigma, SRP0122) incubated with 1 µg of recombinant Oct4 (abcam, ab134876) and Glutathione sepharose 4B (GE healthcare, 17-0756-01) resin in the binding buffer (50 mM Tris pH 8.5, 50 mM KCl, 5 mM MgCl, 0.5% BSA, and 5% glycerol, complemented with a cocktail of protease inhibitors) overnight at 4°C with gentle agitation. The resin was washed twice with binding buffer and proteins were eluted using the elution buffer (50 mM Tris-HCl, 10 mM reduced glutathione, pH 8) according to the manufacturer’s instructions. Eluate and input were loaded onto a 10% SDS-PAGE gels and blots were probed using anti-Lsd1 (Abcam, ab17721) and anti-Oct4 (Santa Cruz, sc-8628) antibodies.

#### *In vitro* Lsd1 demethylase activity assay

An *in vitro* fluorometric assay was used to detect Lsd1 demethylase activity using an Epigenase^TM^ kit (Epigentek, P-0379) according to the manufacturer’s protocol. 0.25 µM of Lsd1 (Sigma, SRP0122) was used for activity assays together with 0.5 µM (or as indicated) of Oct4 (abcam, ab134876 and ab169842) or BSA (Sigma, A3059) or the catalytic domain of Dnmt3a (*purified in-house*) or 0.1 mM of the Lsd1 inhibitor Tranylcypromine (TCP). Signals were measured using a CLARIOstar plate reader and analyzed using MARs software as described by the manufacturer.

#### Histone demethylation assay

Lsd1 histone demethylation assays were performed as described (Shi et al., 2004). A total of 30 µg of bulk histones (Sigma, H9250) in a histone demethylation buffer (50 mM Tris pH 8.5, 50 mM KCl, 5 mM MgCl, 0.5% BSA, and 5% glycerol) were incubated with 0.25 µM of Lsd1 (Sigma, SRP0122) alone, or increasing concentrations (0.125 µM, 0.25 µM, 0.5 µM, 1 µM) of Oct4 (Abcam, ab134876 and ab169842), or 0.1 µM TCP for 4 hr at 37°C. Lsd1 activity was monitored by Western blot using anti-H3K4me2 antibody (Abcam, ab32356). The membrane was stained by Ponceau S to determine equal loading of the reaction mix.

#### Co-immunoprecipitation

Co-immunoprecipitation experiments was also performed as described (Whyte et al., 2012). Briefly, undifferentiated F9 ECCs and ESCs were washed and harvested in cold 1X PBS. Cellular proteins were extracted using TNEN250 lysis buffer (50 mM Tris pH 7.5, 5 mM EDTA, 250 mM NaCl, 0.1% NP-40) complemented with protease inhibitors at 4°C with rotation for 30 min. Complexes were then immunoprecipitated overnight at 4°C with rotation by incubating the supernatant solution supplemented with two volumes of TNENG (50 mM Tris pH 7.5, 5 mM EDTA, 100 mM NaCl, 0.1% NP-40, 10% glycerol) with Dynabeads® M280 (Life Technologies, 11203D) bound to 5 ug of antibody. Beads were washed with TNEN125 (50 mM Tris pH 7.5, 5 mM EDTA, 125 mM NaCl, 0.1% NP-40) and samples were eluted by boiling for 10 min in Laemmli’s loading buffer containing 100 mM DTT. Western blots were performed with NuPAGE 4-12% Tris-Bis gels. Antibodies used included: Lsd1 (abcam, ab17721), HDAC1 (abcam, ab7028), Mi-2b (abcam, ab72418).

#### Western blot

Western blot analysis was performed using the standard method and the following antibodies and dilutions: anti-Dnmt3a, 1:1000 (Active Motif, 39206), Anti-Lsd1, 1:1000 (abcam, ab17721) and anti-β Actin, 1:1000 (Santa Cruz, sc8628), and anti-Rabbit, 1:10,000 (Jackson Immunoresearch, 111-035-003) or anti-Mouse, 1:10,000 (Jackson Immunoresearch, 115-035-003). Chemiluminescence was performed according to the manufacturer’s protocol (Thermo-Fisher Scientific, 34580).

#### MethylRAD sequencing Analysis

##### Alignment and Quality Control

A total of 258,987,008 single end 1×150 sequencing was performed using a NovaSeq 6000 platform for undifferentiated and day 4 F9 samples. The program FastQC v. 0.11.7 (Andrews, 2010) was used to check data quality pre- and post-quality trimming/adapter removal. Adapters were removed from reads using Trimmomatic v. 0.36 (Bolger et al., 2014). Trimmomatic is a program that removes adapter sequences and trims short Illumina reads based on quality. Cutadapt version 2.2 (Martin, 2011) was used to trim reads further, removing the first two and last two bases of each read. Reads containing greater than 0 N’s were discarded. After trimming, a total of 112,289,569 reads remained in the undifferentiated and day 4 samples. Reads which do not have *Fsp*E1 sites present anywhere in the read were removed using ‘grep’, leaving a total of 71,730,977 reads. Finally, Bowtie2 version 2.3.3 (Langmead et al., 2009, Langmead and Salzberg, 2012) was used to map reads to the ENSEMBL *Mus musculus* reference genome version GRCm38.93. A maximum of 1 mismatch was allowed in read mapping. The mapping rate of reads was 89%, with 64% of the reads mapped to the genome exactly 1 time and included in further analyses.

##### Data preprocessing

Methylated sites were catalogued by iterating through all read sequences to find a matched pattern of methylation (i.e. CCGG, CCAGG, and CCTGG) and recording its location in the genome. The number of reads mapping to each methylated site was recorded and adjusted for substitution, deletion, and insertion accordingly. Sites were also matched with the reference genome for verification. Sites that had less than 5 reads were removed from downstream analysis; counts from duplicate sites between patterns were summed as one site. Python and R scripts used in this analysis are included at www.github.com/natallah.

##### Annotation to LSD1 Enhancers

Sites in the LSD1 enhancers were modified to include 1 kb up- and downstream of the identified start site. Both the undifferentiated and day 4 differentiated F9 samples were annotated to the modified LSD1 regions using BEDTools intersect. The total amount of methylation in a region was determined by combining the read counts of all sites in that region. Upper and lower quartiles were used in thresholding regions as gaining or losing methylation. Specifically, all regions with at least 22 counts more in undifferentiated than in day 4 differentiated samples were identified as losing DNA methylation as differentiation occurred. All regions with at least 30 counts more in day 4 samples than in undifferentiated samples were identified as gaining DNA methylation.

##### Quantifying methylated regions

The union of enhancers found in both the undifferentiated and day 4 differentiated F9 samples were determined. The difference in methylation for each combined enhancer regions were computed by subtracting the total methylation in day 4 differentiated F9 samples to that of the undifferentiated sample. Comparative figures were produced from these final data.

##### Determining methylation level of enhancers

DNA sequences of all known enhancers for the ESC_J1 strain of mouse were downloaded from EnhancerAtlas V2.0 (Gao et al., 2016). Overlaps between MethylRAD sites and enhancers were found using BEDTools intersect. Reads that overlapped the enhancer for the same gene were then summed together. Raw counts were normalized for length. The average length of enhancers in the EnhancerAtlast database was computed for each gene. If different studies in the database reported variable lengths for enhancers, the average length of the enhancer was computed. Counts for each sample were then divided by the length of the enhancer and subsequently multiplied by 1000 (to enhance readability). The 25^th^ and 75^th^ percentiles were computed for all enhancers separately for undifferentiated and day four samples and were used as cutoffs for low and intermediate methylation. Thus, for undifferentiated samples, enhancers annotated as having low methylation have normalized counts between (0, 24.57], intermediate methylation are between (24.57, 243.68], and high methylation have greater than 243.58 normalized counts. Enhancers in day 4 samples are annotated as highly methylated if normalized counts are observed to be between (0, 24.06], intermediate counts are between (24.06, 256.56], and high counts have greater than 256.56 normalized counts.

#### ChIP-Seq Analysis

##### Quality Control and Mapping

Sequencing was performed using a NovaSeq 6000 to generate > 80 million paired-end (2×50) reads (>80 million) undifferentiated and day 4 F9 samples. Sequence data quality was determined using FastQC software (Andrews, 2010) and quality based trimming and filtering (minimum quality score 30 and minimum read-length 20) was performed through TrimGalore tool (Krueger, 2012). Greater than 95% of the reads from all samples were retained after quality control and were used for the mapping. Mapping was performed against the mouse reference genome (GRCm38) using Bowtie2 (Langmead and Salzberg, 2012) with a maximum of 1 mismatch. The overall mapping rate was >97% for all samples. Bowtie2 derived BAM files were further filtered to retain the reads with minimum MAPping Quality (MAPQ) 10.

##### Peak-calling

Peak calling was performed using epic2 (Stovner and Saetrom, 2019) for each Input-ChIP pair using the mouse reference genome (GRCm38). The tool was run with MAPQ10 filtered BAM files and the default parameters (--falsediscovery-rate-cutoff 0.05, --binsize 200).

##### Peak-annotation

Peak-annotation and visualization was performed with R-package ChIPseeker (Yu et al., 2015a). Annotations were performed for all epic2 peaks called with default parameters.

##### Overlap of H3K4me1 peaks in F9 ECCs with those in ESCs

Overlapping LSD1 bound sites between H3K4me1 peaks in F9 ECCs and ESCs (Whyte et al., 2012). Correct overlap, was determined by converting peak coordinates from mm9 to mm10 using CrossMap tool (Zhao et al., 2014).

#### Pathway Analysis

IPA (Ingenuity Pathway Analysis), (IPA, QIAGEN Redwood City, www.qiagen.com/ingenuity), was used in the annotation of genes and in performing the pathway analyses.

### QUANTIFICATION AND STATISTICAL ANALYSIS

#### methylRAD statistical analyses and software

-Methods are described in the section entitled “DNA methylation is not established at PpGe during F9 ECC differentiation”, and also in the Method Details section entitled “MethylRAD sequencing Analysis.”

-fastQC version 0.11.7 was used to check data quality before and after filtering.

-Read trimming was performed using Trimmomatic v. 0.36 to remove Illumina adapter sequences from reads. Reads containing greater than 0 N’s were discarded.

-Cutadapt v 2.2 was used to remove the first two and last two bases of each read.

-grep “CCGG|GGCC|CCAGG|GGTCC|CCTGG” *fastq > sites.fastq was used to keep only reads with FspEI sites present

-Bowtie v2.3.3 was used to map reads to the GRCm38.93 Mus Muscusus genome, removing all reads with more than 1 mismatch

-Methylated sites were identified and methylation events counted using a custom Python script, which is available at www.github.com/natallah

-A total of 1,370,254 cytosines were captured genome-wide with a cutoff of 5 reads per site minimum (1,370,254 cytosines had at least 5 reads aligned with Bowtie2). The difference in methylation was calculated by summing reads which overlap with enhancer downloaded from EnhancerAtlas 2.0.

-Differences were calculated between differentiated and undifferentiated samples by calculating the difference d = xij-xij with x equivalent to the read count (total raw reads) for enhancer i in sample j. Undifferentiated enhancer read counts were subtracted from differentiated enhancer counts. Special focus was given to the enhancers which were identified as Lsd1 bound previously (Whyte et al, 2012). Low-intermediate-high levels of methylation were calculated using the total read counts in all enhancers from the EnhancerAtlas by using the 75th percentile as the lower bound for high methylation levels and the 25th percentile as the upper bound for low methylation levels.

-BEDtools overlap was used to identify reads overlapping with specific genomic features (enhancers, Lsd1 bound enhancers, promotors)

-ChIPseeker v 1.22.0 was used to annotate site distribution across the genome.

#### ChIP-seq statistical analyses and software

-Methods are described in the section entitled “High throughput analysis of changes in H3K4me1 at PpGe” as well as in the Supplemental section entitled “ChIP-Seq Analysis”.

-FastQC v 0.11.7 was used to check read quality before and after read trimming.

-Bowtie2 v 2.3.3 was used to map to the mouse reference genome version GRCm38, removing alignments with greater than 1 mismatch. Reads with less than a MAPQ of 10 were removed.

-TrimGalore v 0.6.4 was used to trim adapter sequences and remove reads with under a read-length of 20 and with less than a quality PHRED score of 30.

-Peak calling was performed using Epic2 v 2019-Jan-03 with all defaults on each IP and input pair. The peaks were filtered selecting only peaks with false discovery rate less than or equal to 5% and with greater than or equal to a log fold-change (IP/input control) of 2. --binsize was set to 200 in epic2.

-Peaks which had greater than a z-score of 1 were counted as increased, while those with below a z-score of −1 were counted as decreased. The peaks with a z-score greater than or equal to −1 and less than or equal to 1 were annotated as not changing (see Figure 4). Z-score was calculated as how many standard deviations a certain value is above or below the mean using formulae z = (x – μ) / σ, where x corresponds to difference between log2FC of D4 and undifferentiated samples for a specific gene, μ is mean difference across all genes, and σ is standard deviation of difference across all genes.

#### Pathway analyses

-All pathway analyses were performed using Ingenuity Pathway analysis (IPA), as described in the section entitled “DNA methylation is not established at PpGe during F9 ECC differentiation”, as well as the Supplemental section entitled “Pathway Analysis” using Mus musculus as the organism, and input as genes which are low (less than or equal to the 25th percentile methylation), intermediate (greater than the 25th percentile and less than the 75th percentile for all enhancers from EnhancerAtas 2.0), or high (greater than or equal to 75th percentile methylation).

-A one-tailed Fisher’s exact test was used to calculate functional enrichment, and all p-values were adjusted for multiple testing using the Benjamini-Hochberg method. The adjusted p-value cutoff for significance is padj <0.05.

-ggplot was used to generate pie charts, waterfall plots, and bar charts.

### DATA AND CODE AVAILABILITY

-The datasets generated during this study are available at GEO under accession <u>GSE135225</u> (ChIP-seq) and GSE135226 (methylRAD-seq). ChIP-seq peak annotations and genome browser track files are included.

-All code and scripts use to analyze data and to generate genomic plots are at www.github.com/natallah

**Table S4. A list of all PCR primers used in this study. Related to DNA methylation analysis, Chromatin Immunoprecipitation and ChIP-Seq and Gene expression analysis in Method Details**

A list of all PCR primers used in this study (5’ to 3’), separated by technique. The presence of an (AminoC6) modification at the 3’ end of the antisense oligonucleotide of each adaptor is used to block extension

## KEY RESOURCES TABLE

| REAGENT<br>or<br>RESOURCE | SOURCE | IDENTIFIER |
| --- | --- | --- |
| Antibodies |  |  |
| anti-cMyc antibody | Millipore | MABE282 |
| anti-SSEA-1 | Millipore | MAB430 |
| and Alexa Fluor 555 nm | Life Technologies | A21422 |
| anti-Oct4 | Abcam | ab181557 |
| anti-CHD4 | Abcam | ab72418 |
| anti-HDAC1 | Abcam | ab7028 |
| anti-Lsd1 | Abcam | ab17721 |
| anti-HBO1 | Abcam | ab124993 |
| anti-Oct4 | Santa Cruz | sc-8628 |
| anti-H3K4me2 | Abcam | ab32356 |
| anti-H3K27Ac | Abcam | ab4729 |
| anti-H3K4me1 | Abcam | ab8895 |
| anti-H3K27me3 | Abcam | ab8898 |
| anti-β Actin | Santa Cruz | sc8628c |
| Anti-Rabbit IgG | Jackson ImmunoResearch | 111-035-003 |
| Anti-Mouse IgG | Jackson ImmunoResearch | 115-035-003 |
| Biological Samples |  |  |
| pCAG-Myc-Oct4 | Addgene | 13460 |
| Chemicals, Peptides, and Recombinant Proteins |  |  |
| TRIzol | Invitrogen | 15596026 |
| FspEI | NEB |  |
| Protein A magnetic beads | Life Technologies | 10002D |
| Protein G magnetic beads | Life Technologies | 10004D |
| RNase | Roche | 11119915001 |
| Proteinase K | Worthington | LS004222 |
| Quick-RNA<sup>T</sup><sub>M</sub> MiniPrep Plus Kit | ZymoResearch | R1057 |
| Verso One-Step RT-qPCR kit | Thermo Scientific | AB-4104A |
| Alkaline phosphatase staining kit | Sigma | AB0300 |
| GST-Lsd1 | Sigma | SRP0122 |
| Oct4 | Abcam | ab134876 |
| Oct4 | Abcam | ab169842 |
| Bulk Histones | Sigma | H9250 |
| EpiTe<sup>ct</sup> Fast Bisulfite Conversion Kit | Qiagen | 59802 |
| Lipofectamine 2000 | ThermoFischer | 11668019 |
| Critical Commercial Assays |  |  |
| Epigenase<sup>T</sup> M kit | Epigentek | P-0379 |
| Deposited Data |  |  |
| Raw and analyzed data | This paper | GSE135225 |
| Experimental Models: Cell Lines |  |  |
| P19 embryonal carcinoma cells | ATCC | CRL-182 |
| F9 embryonal carcinoma cells | ATCC | CRL-1720 |
| E14Tg2A Embryonic stem cells | MMRRC | 015890-UCD |
| Oligonucleotides |  |  |
| List of primers used | This paper | Table S4 |
| Software and Algorithms |  |  |
| Trimmomatic v 0.36 | Bolger, A. M., Lohse, M., & Usadel, B. (2014). Trimmomatic: A flexible trimmer for Illumina Sequence Data. *Bioinformatics*, btu170. | https://github.com/timflutre/trimmomatic |
| Bowtie2 v.2.3.3 | Langmead B, Salzberg S. Fast gapped-read alignment with Bowtie 2. *Nature Methods*. 2012, 9:357-359. | http://bowtie-bio.sourceforge.net/bowtie2/manual.shtml |
| Cutadapt v 2.2 | MARTIN, Marcel. Cutadapt removes adapter sequences from high-throughput sequencing reads. **EMBnet.journal**, [S.l.], v. 17, n. 1, p. pp. 10-12, may 2011. ISSN 2226-6089. Available at: < http://journal.embnet.org/index.php/embnetjournal/article/view/200 >. Date accessed: 11 dec. 2019. doi: https://doi.org/10.14806/ej.17.1.200. | https://cutadapt.readthedocs.io/en/stable/installation.html |
| ENSEMBL Mus musculus reference genome version GRCm38.93 | GRCm38.p2 (Genome Reference Consortium Mouse Reference 38), INSDC Assembly GCA_000001635.4, Jan 2012 | ftp://ftp.ensembl.org/pub/release-98/fasta/mus_musculus/dna/ |
| Custom python scripts | https://github.com/natallah/methylRad | https://github.com/natallah/methylRad |
| Bismark v 0.18.2 | Felix Krueger, Simon R. Andrews, Bismark: a flexible aligner and methylation caller for Bisulfite-Seq applications, *Bioinformatics*, Volume 27, Issue 11, 1 June 2011, Pages 1571–1572, https://doi.org/10.1093/bioinformatics/btr167 | https://www.bioinformatics.babraham.ac.uk/projects/bismark/ |
| BEDTools v 2.29.0 | Quinlan AR, Hall IM. BEDTools: a flexible suite of utilities for comparing genomic | https://bedtools.readthedocs.io/en/latest/ |
|  | features. *Bioinformatics*. 2010;26(6):841–842. doi:10.1093/bioinformatics/btq033 |  |
| Enhancer Atlas v 2.0 | Tianshun Gao and Jiang Qian. *EnhancerAtlas 2.0: an updated resource with enhancer annotation in 586 tissue/cell types across nine species*. *Nucleic Acids Res*. 2019. | http://enhanceratlas.org/ |
| TrimGalore v 0.6.4 | Krueger F: Trim Galore!. [http://www.bioinformatics.babraham.ac.uk/projects/trim_galore/] | https://www.bioinformatics.babraham.ac.uk/projects/trim_galore/ |
| epic2 2019-Jan-03 | Endre Bakken Stovner, Pål Sætrom, epic2 efficiently finds diffuse domains in ChIP-seq data, *Bioinformatics*, Volume 35, Issue 21, 1 November 2019, Pages 4392–4393, https://doi.org/10.1093/bioinformatics/btz232 | https://bioepic.readthedocs.io/en/latest/ |
| FastQC v. 0.11.7 | Andrews S. (2010). FastQC: a quality control tool for high throughput sequence data. Available online at: http://www.bioinformatics.babraham.ac.uk/projects/fastqc | https://www.bioinformatics.babraham.ac.uk/projects/fastqc/ |
| ChIPseeker v 1.22.0 | Yu G, Wang L, He Q (2015). “ChIPseeker: an R/Bioconductor package for ChIP peak annotation, comparison and visualization.” *Bioinformatics*, **31** (14), 2382-2383. doi: [10.1093/bioinformatics/btv145](https://doi.org/10.1093/bioinformatics/btv145). | https://bioconductor.org/packages/release/bioc/html/ChIPseeker.html |
| Ingenuity Pathway Analysis (IPA) | Qiagen | https://www.qiagenbioinformatics.com/products/ingenuity-pathway-analysis/?gclid=Cj0KCQiAuefvBRDXARIsAFEOQ9GUaRrR3QZy42IJtb5QHvIHsmCrtw5fOeyRo0d2GEgfWxKTIPwd8w8aArVwEALw_wcB |

---

**Table S4.**
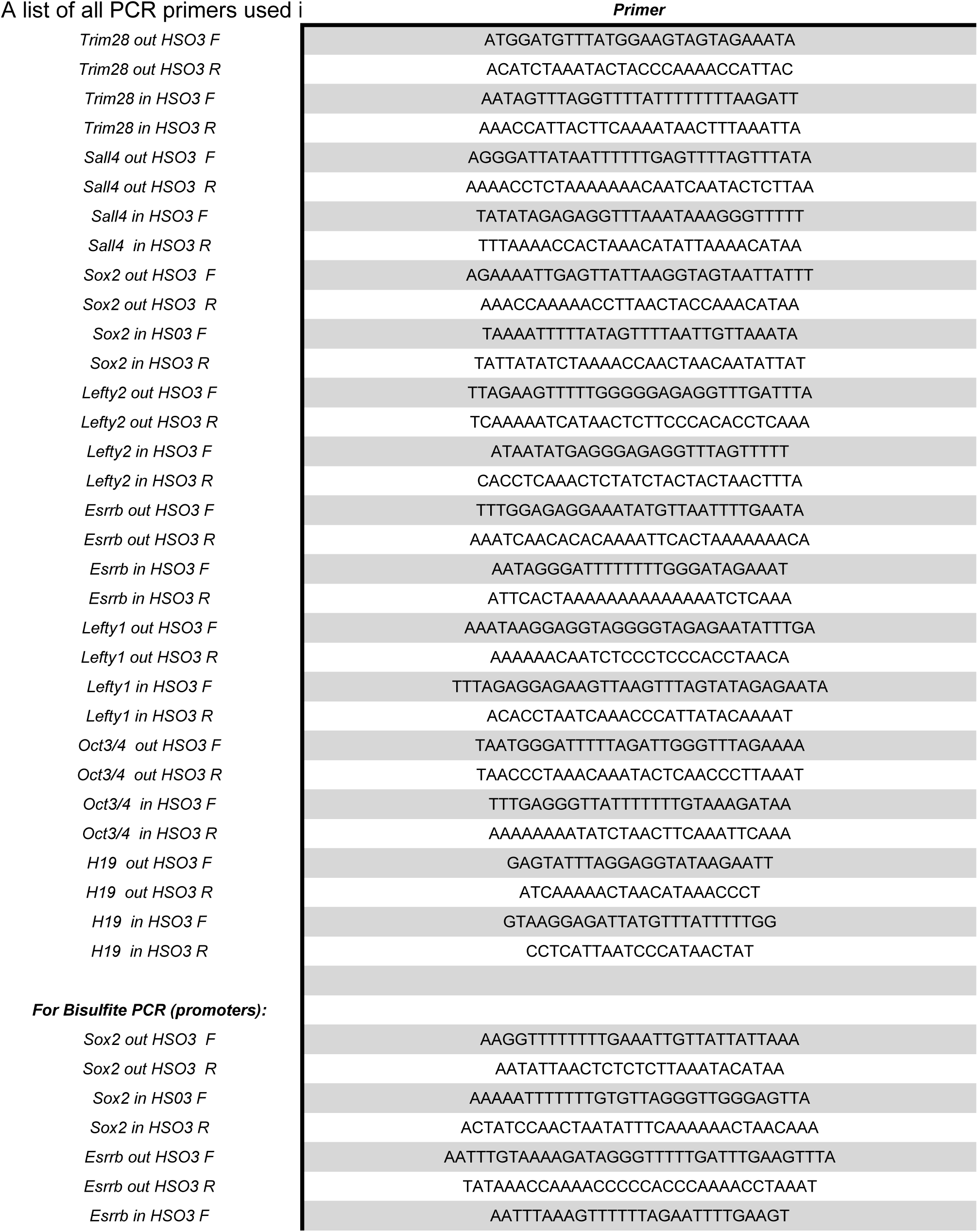
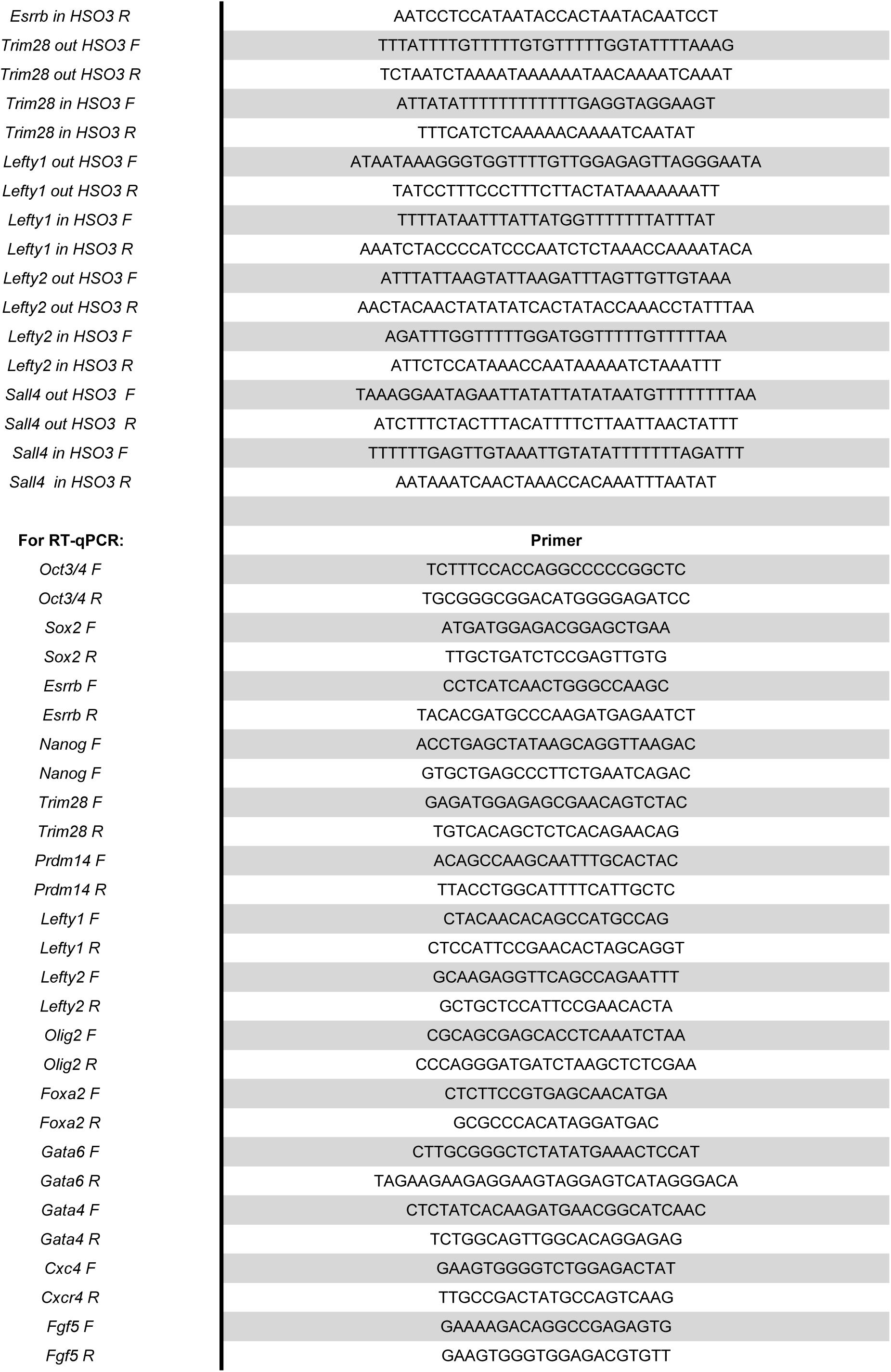
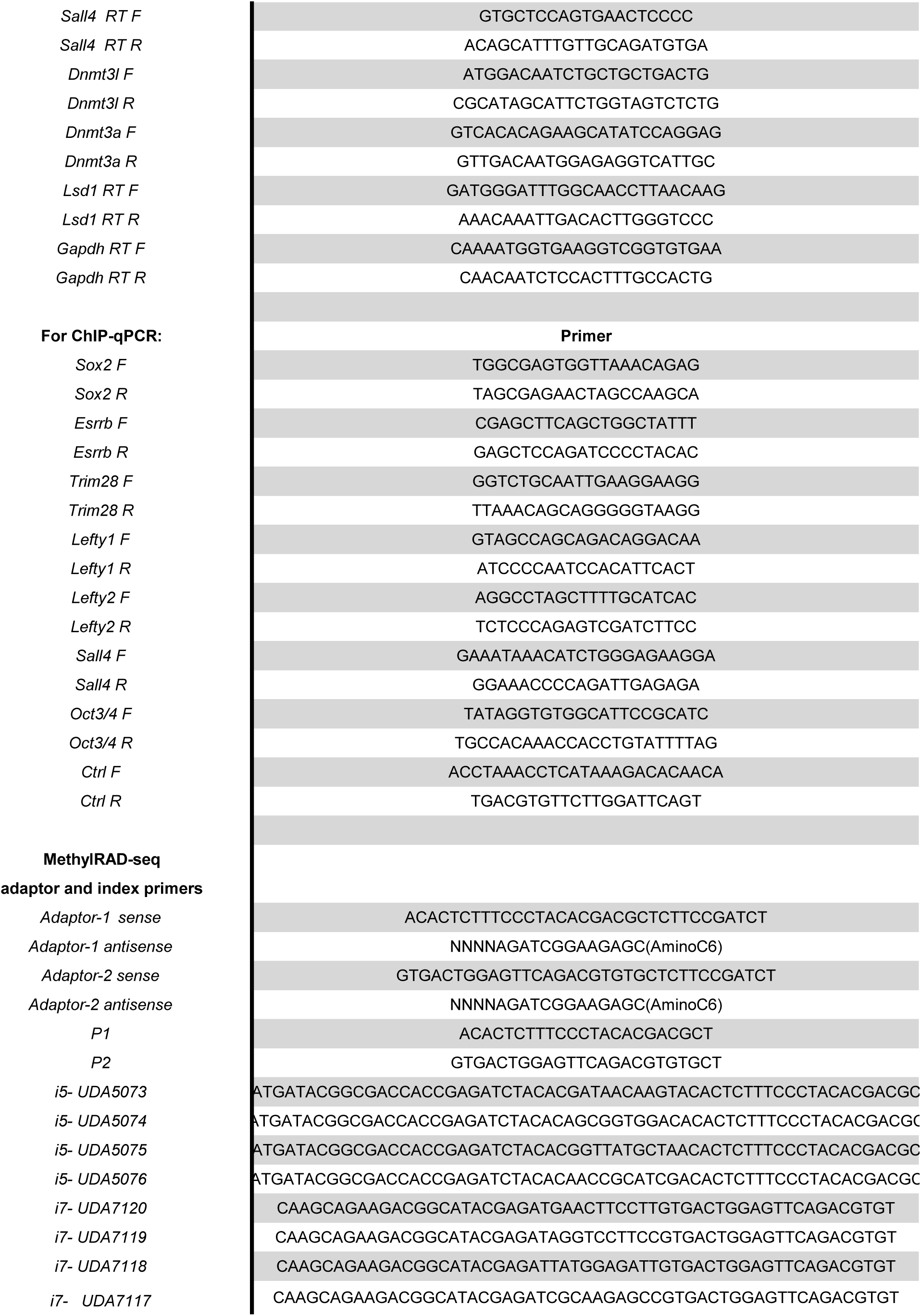
Primers used in this study.

**Figure S1.**
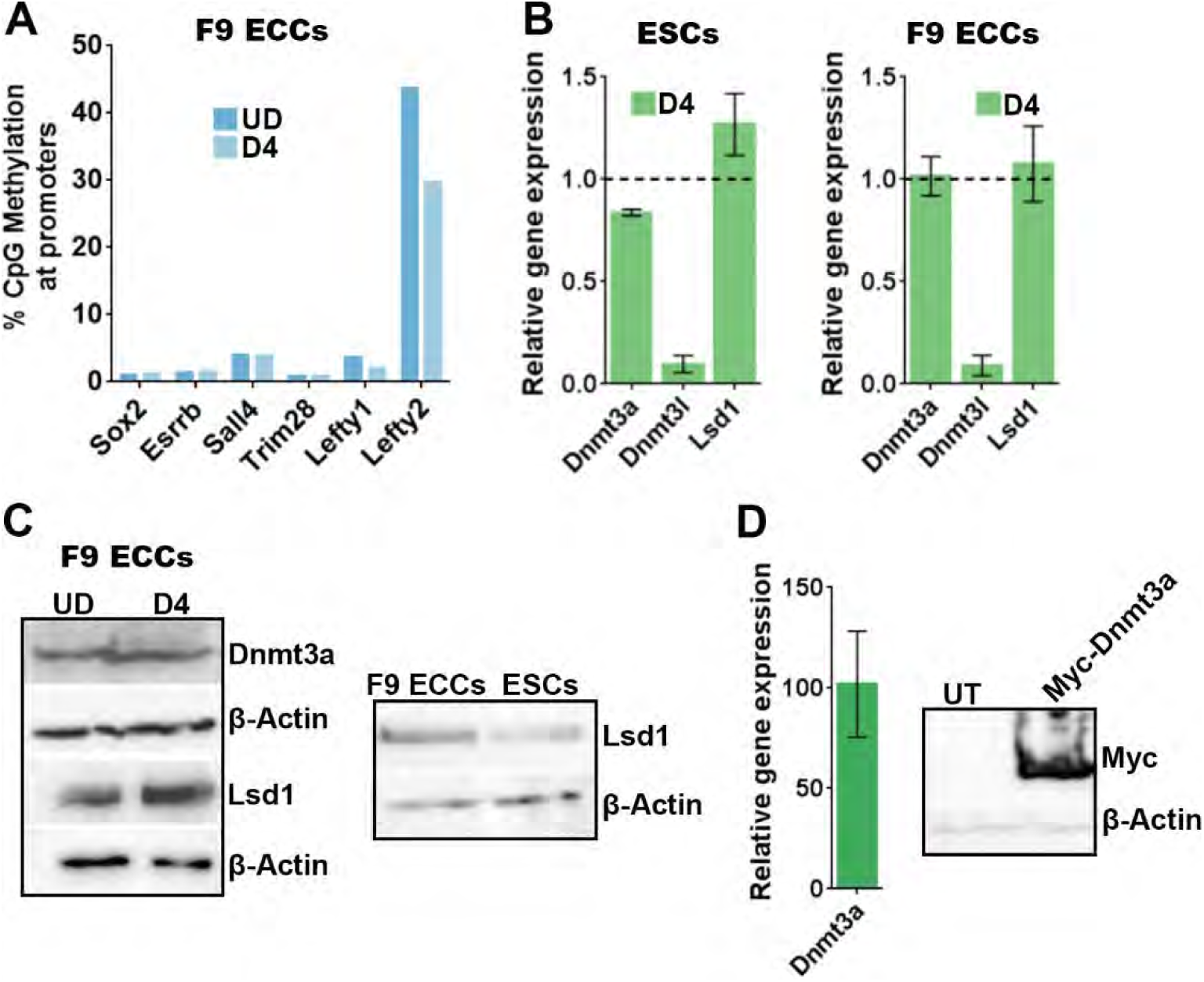
Expression of epigenetic effectors in embryonal carcinoma cells. (Related to Figure 1) UD: undifferentiated, D4: Days post-differentiation. ESCs: embryonic stem cells and F9 ECCs: F9 embryonal carcinoma cells; PpGe: pluripotency gene enhancers. (A) Bis-Seq analysis of DNA methylation at PpG promoters in F9 ECCs pre- and post-differentiation. DNA methylation at PpG promoters remained under 10% except *Lefty2*, which shows very high methylation in the UD state that is reduced post-differentiation. (B) RT-qPCR comparing the changes in expression of DNA methyltransferase Dnmt3a and histone demethylase Lsd1 in ESCs with that in F9 ECCs pre- and post-differentiation. The C_t_ values are normalized to *Gapdh* and represented relative to expression in undifferentiated cells (dotted line). In both ESCs and ECCs, Lsd1 and Dnmt3a expression is maintained post-differentiation. (C) Western blot. A total of 50 µg of total protein from undifferentiated and differentiated cells was loaded in each well. Left panel confirms comparable expression of Dnmt3a and Lsd1 pre- and post-differentiation in F9 ECCs. Right panel compares Lsd1 expression in F9 ECCs with ESCs showing very similar levels in these cells. β-Actin is the loading control. (D) Gene expression analysis by RT-qPCR and Western blot confirming recombinant Myc-Dnmt3a overexpression in F9 ECCs 24 hr post-differentiation (48 hr post-transfection). The data are normalized to a *Gapdh* control and shown relative to untransfected that is set to 1. β-Actin is used as loading control for Western blot.

**Figure S2.**
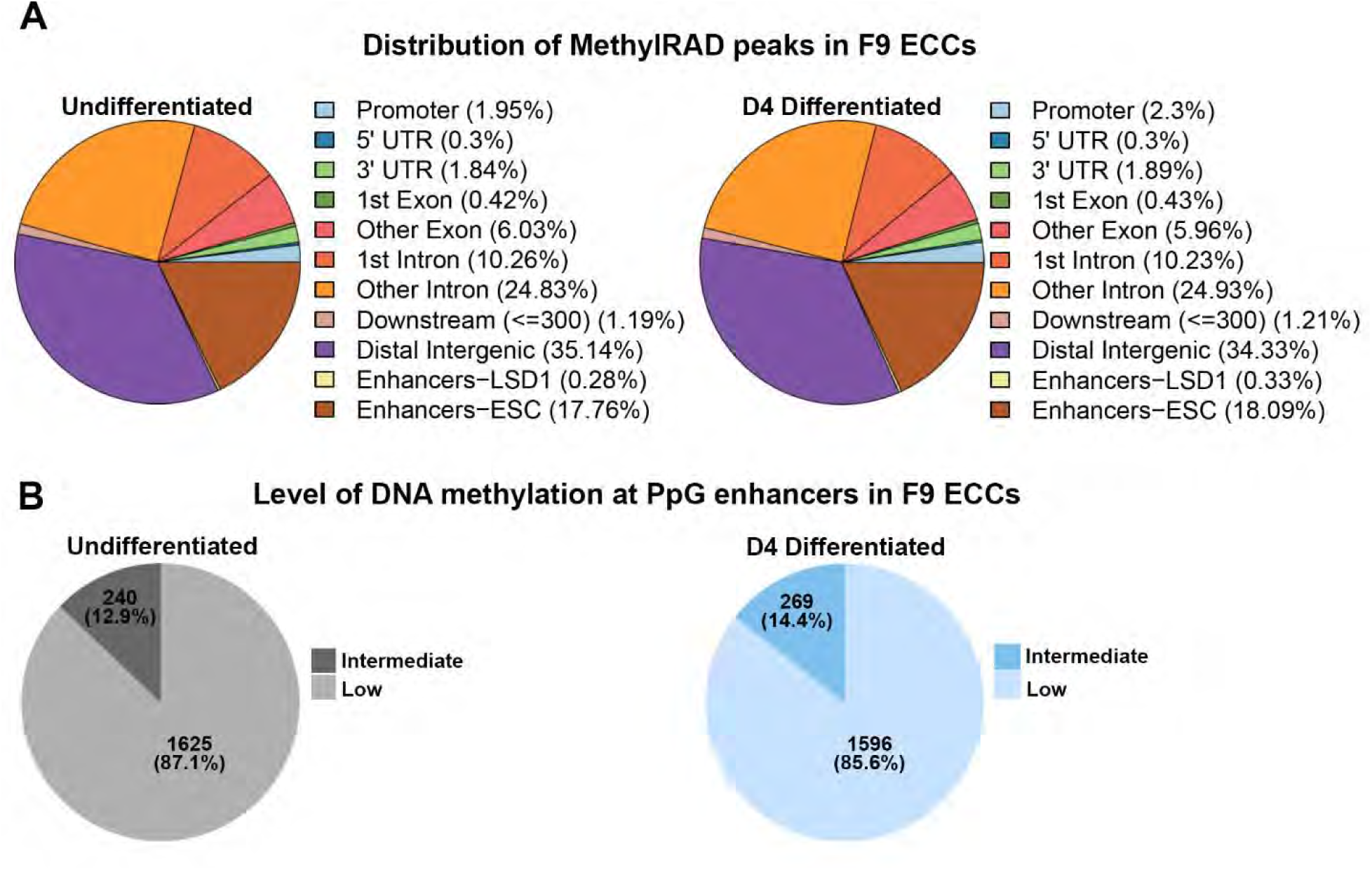
Distribution of MethylRAD peaks. (Related to Figure 2) (A) Fractional distribution of MethylRAD peaks in undifferentiated and differentiated F9 ECCs across regulatory regions of the genome. (B) Pie charts show the level of methylation at PpGe in undifferentiated (grey) and differentiated day 4 (blue) samples. Most PpGe had low levels of methylation or to a lesser extent, intermediate levels of methylation, with none having high CpG methylation.

**Figure S3.**
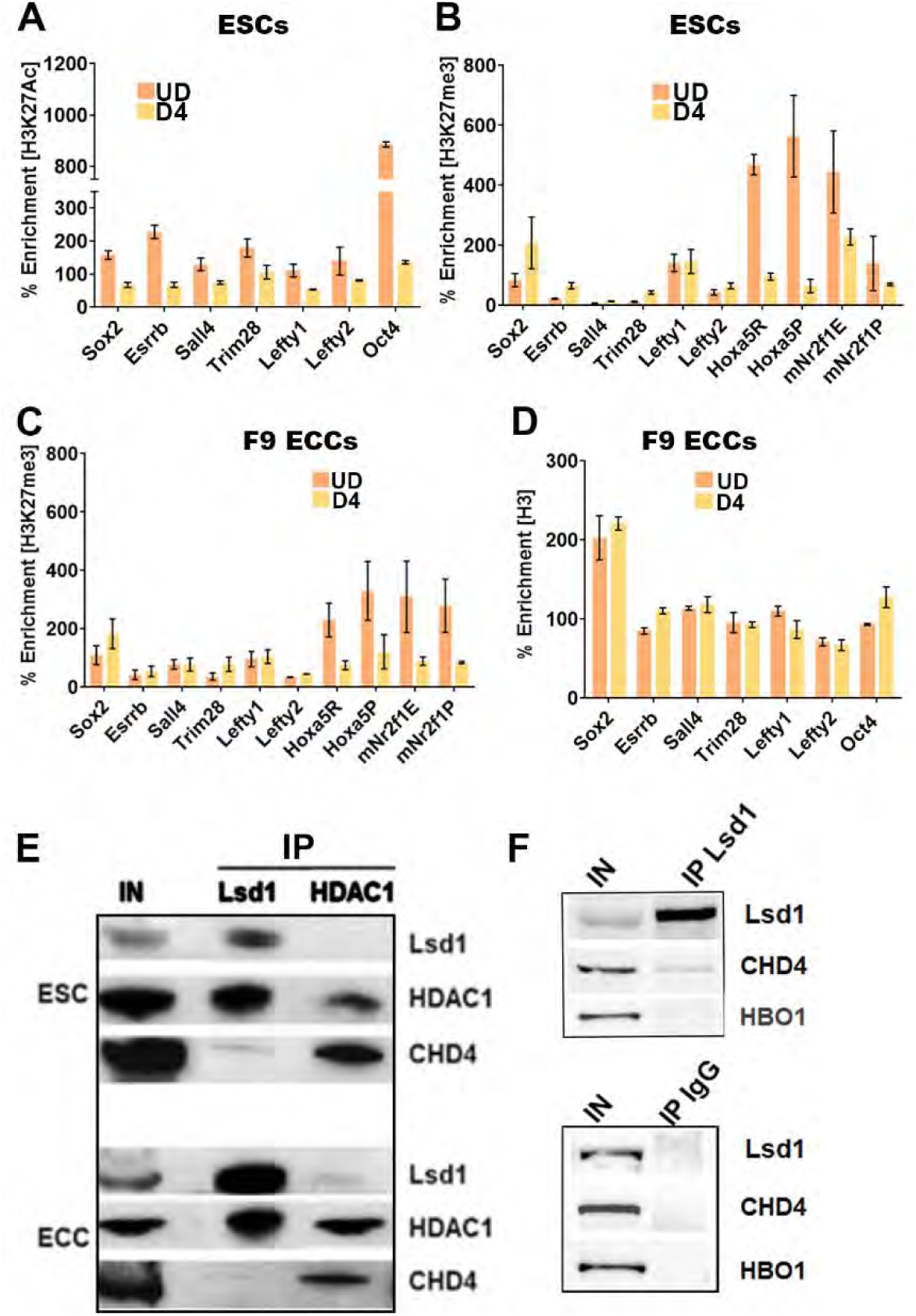
H3K27 modification of pluripotency gene enhancers in embryonic stem cells and embryonal carcinoma cells. (Related to Figure 3) UD: undifferentiated; D4: Days post-induction of differentiation; IP: immunoprecipitation (A, B, C and D) ChIP-qPCR was used to determine the enrichment of histone modifications at PpGe. (A) H3K27Ac in ESCs (B) H3K27me3 in ESCs (C) H3K27me3 and (D) H3 in F9 ECCs pre- and post-differentiation. Whereas deacetylation of PpGe is observed as a decrease in H3K27Ac signal post-differentiation in ESCs, there is no gain of H3K27me3 at these sites, neither in ESCs nor in F9 ECCs. As previously reported, we observed a decrease in H3K27me3 at the enhancers and promoters of *Hoxa5* and *mNr2f1* genes, consistent with their transcriptional activation in response to differentiation (Laursen, Mongan et al. 2013). No change was observed in H3 occupancy between UD and D4 at PpGe. % Enrichment = Fold enrichment over input X100 (E, F) Co-IP was performed with anti-Lsd1 or anti-HDAC1 and a control IgG using whole cell extracts from undifferentiated ESCs and F9 ECCs. 20% of the input and eluate from Co-IP were probed for Lsd1-Mi2/NuRD subunits (Lsd1, HDAC1 and CHD4) on Western blot and HBO1 was used as a negative control.

**Figure S4.**
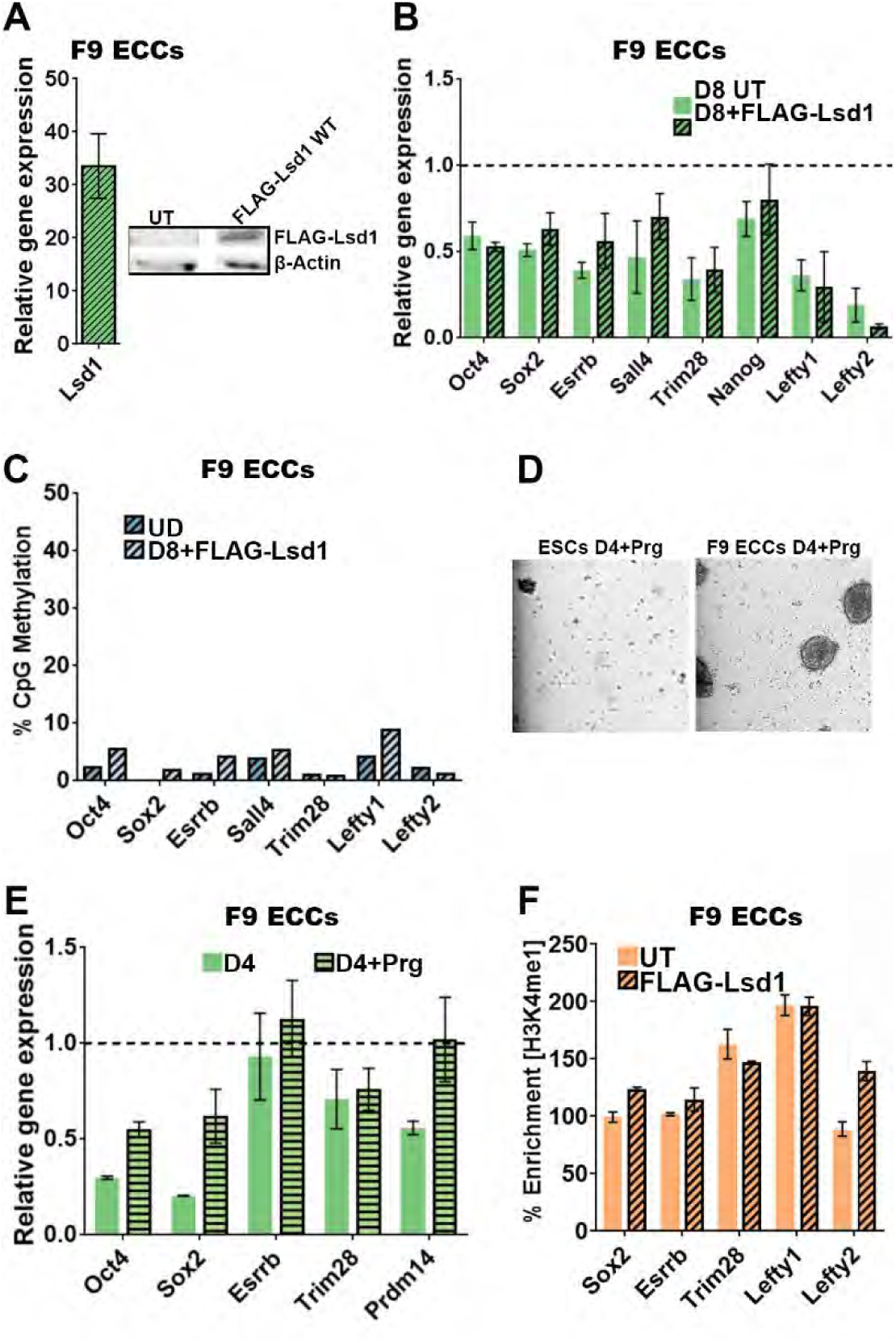
Effect on Lsd1 on differentiation and pluripotency gene enhancer silencing in embryonal carcinoma cells. (Related to Figure 3) UD: undifferentiated; D8: Days post-induction of differentiation; Prg: pargyline. (A) Gene expression analysis by RT-qPCR and Western blot examining the expression of recombinant FLAG-Lsd1 in F9 ECCs 24 hr post-differentiation (48 hr post-transfection). The C_t_ values are normalized to *Gapdh* and represented relative to expression in untransfected cells (set to 1). β-Actin is used as loading control for Western blot. (B) Gene expression analysis by RT-qPCR of PpGs in F9 ECCs expressing FLAG-Lsd1. Similar to our observation in untransfected WT F9 ECCs, PpGs are partially repressed in these cells, showing no effect of recombinant Lsd1 on PpG repression (*p-*value >0.1). (C) Bis-Seq analysis of DNA methylation at PpGe in F9 ECCs expressing recombinant FLAG-Lsd1 showed no gain, corresponding to no loss of H3K4me1 at these sites. Data are an average and SEM of two biological replicates. (D) Bright field microscopy of ESCs and ECCs differentiated for 4 days in the presence of the Lsd1 inhibitors Prg. 80-90% cell death is observed in ESCs. Lsd1 inhibitor has no effect on F9 ECC differentiation as shown by a normal morphology of embryoid bodies. Scale bar is 100 µm. (E) Gene expression analysis of PpGs by RT-qPCR in differentiating F9 ECCs untreated or treated with Prg. A slight derepression of some PpGs was observed in inhibitor treated cells post-differentiation. Gene expression was normalized to *Gapdh* and represented as relative change to gene expression in undifferentiated (dotted line). (F) Undifferentiated F9 ECCs were transfected with FLAG-Lsd1 and cultured for 72 hr. H3K4me1 enrichment was determined using ChIP-qPCR. No change in H3K4me1 levels was observed compared to untransfected (UT) cells. % Enrichment = Fold enrichment over input X100

**Figure S5.**
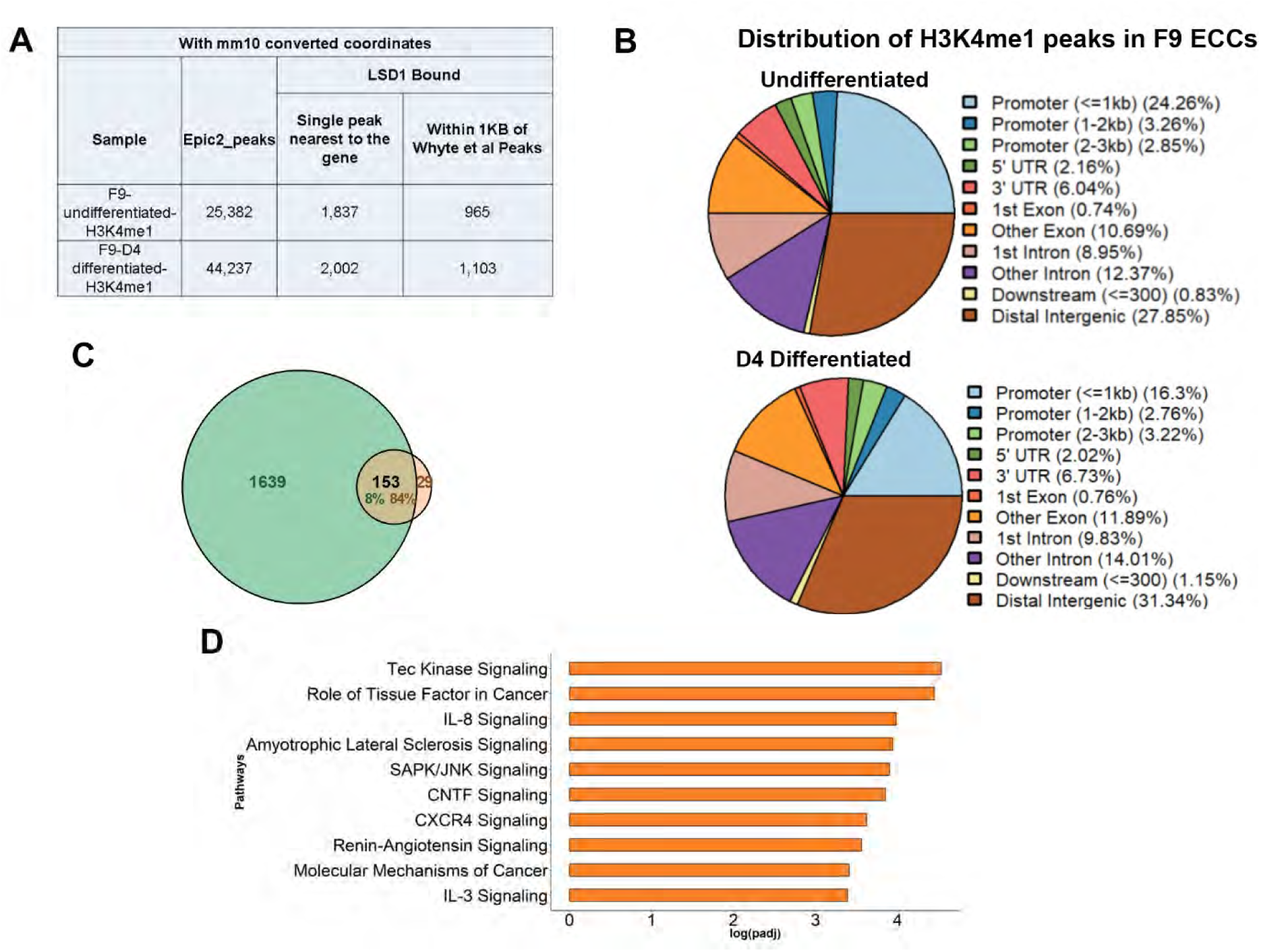
Distribution of H3K4me1 ChIP-Seq peaks. (Related to Figure 4) (A) Summary of the overlapping LSD1 bound sites between H3K4me1 peaks in F9 ECCs and ESCs. To determine correct overlap, peak coordinates from Whyte.et al were converted from mm9 to mm10 using CrossMap tool. (B) Fractional distribution of H3K4me1 peaks in undifferentiated and differentiated F9 ECCs throughout the genome. (C) Overlap of the 182 sites that show decrease in H3K4me1 post-differentiation in F9 ECCs with those in ESCs (See Supplementary methods). (D) Top ten most statistically significant enriched canonical pathways amongst the genes associated with decrease in F9 ECCs.

**Figure S6.**
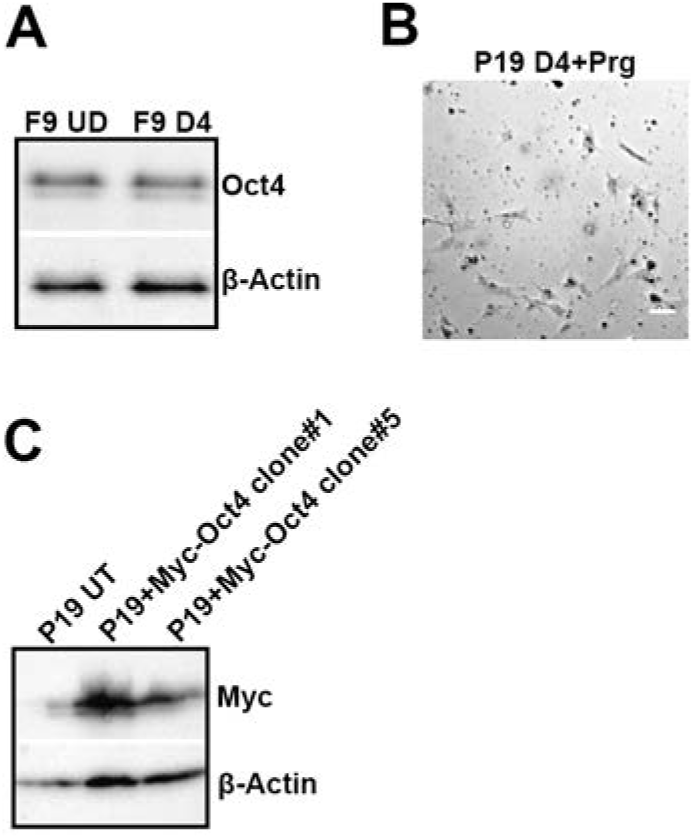
P19 embryonal carcinoma cells are sensitive to Lsd1 inhibition. Related to Figure 5 and Figure 6. D4: Days post-induction of differentiation; Prg: pargyline; UT: untransfected control. (A) Western blot showing protein levels of Oct4 in undifferentiated and D4 differentiated F9 ECCs. A continued expression of Oct4 is seen in differentiated cells although it is lower than that observed in the undifferentiated cells. β-Actin was used as a loading control. (B) Bright field microscopy of P19 ECCs differentiated for 4 days in presence of Lsd1 inhibitor, Prg. 80-90% cell death is observed in P19 ECCs indicating that Lsd1 activity at PpGe is required for differentiation. Scale bar is 100 µm. (C) Expression analysis by Western blot confirming recombinant Myc-Oct4 overexpression in two independent clones of P19 ECCs. β-Actin was used as a loading control for Western blot.

## Supplementary Tables

**Table S1.**
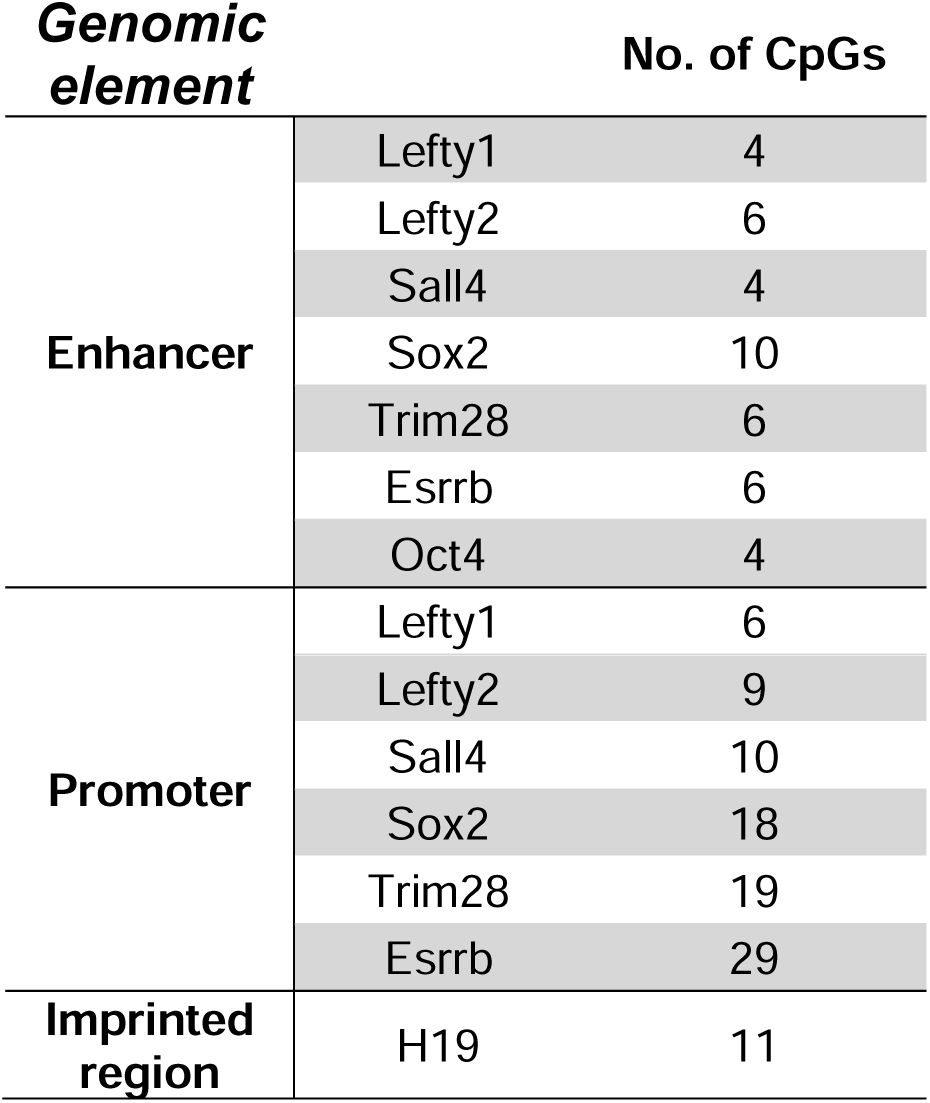
Number of CpGs at each site used for Bis-Seq DNA methylation analysis. Related to DNA methylation analysis in Method Details. The number of CpG sites used to compute percent methylation within the H19 imprinted region, PpG enhancers, and promoters.

**Table S2.** Number of reads for each enhancer used for Bis-Seq analysis. Related to DNA methylation analysis in Method Details. The total number of reads from Wide-Seq runs that were used for data presented in this study. The number of reads were calculated for each sample and enhancer, along with the overall total number of reads.

| *Sample* | *Lefty1* | *Lefty2* | *Sall4* | *Sox2* | *Trim28* | *Esrrb* | *Oct4* | ***Total*** |
| --- | --- | --- | --- | --- | --- | --- | --- | --- |
| *ESCs UD* | 28465 | 21981 | 2861 | 5826 | 17973 | 6034 | 1902 | **85042** |
| *ESCs D4* | 401 | 275 | 216 | 881 | 269 | 5381 | 6322 | **13745** |
| *F9 ECCs UD* | 57536 | 49939 | 27259 | 40357 | 14261 | 22395 | 6322 | **232670** |
| *F9 ECCs D4* | 62876 | 38534 | 29204 | 32570 | 13304 | 21569 | 8865 | **214866** |
| *F9 ECCs D6* | 62185 | 38590 | 30187 | 28239 | 10163 | 13059 | 6980 | **182422** |
| *F9 ECCs+Myc-Dnmt3a UD* | 3733 | 2692 | 1308 | 2612 | 1249 | 24671 | 311 | **38969** |
| *F9 ECCs+Myc-Dnmt3a D8* | 2734 | 1637 | 1135 | 2665 | 1097 | 4316 | 350 | **24705** |
| *F9 ECCs+FLAG-Lsd1 UD* | 2513 | 2267 | 816 | 4204 | 820 | 24412 | 1791 | **43102** |
| *F9 ECCs+FLAG-Lsd1 D8* | 2819 | 1950 | 1946 | 4530 | 1169 | 21576 | 1601 | **42022** |
| *P19 ECCs UD* | 26651 | 18427 | 10541 | 31377 | 9437 | 46154 | 7044 | **175098** |
| *P19 ECCs D4* | 40376 | 30470 | 18198 | 27196 | 11267 | 56736 | 14506 | **241010** |
| *P19 ECCs D8* | 4713 | 8696 | 5880 | 28944 | 6475 | 24128 | 4498 | **96640** |
| *P19 ECCs+Myc-Oct4 UD* | 23029 | 11988 | 11538 | 9324 | 7735 | 28746 | 7510 | **130029** |
| *P19 ECCs+Myc-Oct4 D4* | 35014 | 20519 | 14300 | 18022 | 13227 | 43559 | 14989 | **210333** |
| ***Total*** | **353045** | **247965** | **156838** | **236746** | **108446** | **342735** | **76669** | **1522445** |

**Table S3.** Number of reads for promoters and imprinted regions used for Bis-Seq analysis. Related to DNA methylation analysis in Method Details. The total number of reads from Wide-Seq runs that were used in this study. The number of reads were calculated for each sample, promoter site, and imprinted locus, as well as the overall total number of reads.

| *Sample* | *Lefty1* | *Lefty2* | *Sall4* | *Sox2* | *Trim28* | *Esrrb* | *H19* | ***Total*** |
| --- | --- | --- | --- | --- | --- | --- | --- | --- |
| *F9 ECCs UD* | 51 | 726 | 486 | 9556 | 7575 | 4191 | 11062 | **33647** |
| *F9 ECCs D4* | 99 | 1590 | 1400 | 17431 | 16307 | 13865 | 9376 | **60068** |
| ***Total*** | **150** | **2316** | **1886** | **26987** | **23882** | **18056** | **20438** | **93715** |

